# YAP disrupts bile acid homeostasis to drive cancer-associated cachexia

**DOI:** 10.64898/2026.02.01.702698

**Authors:** Madeline L. Webb, Mikaela Wong, Anthony P. Karamalakis, Yuchen Liu, Kimberley J. Evason, Kira X. Sun, Kristin K Brown, Jay R. Black, Yingzi Yang, Andrew G. Cox

## Abstract

Cancer-associated cachexia is a severe metabolic syndrome marked by dramatic loss of adipose and muscle mass. Although preclinical models have advanced our understanding of cachexia, there are still no approved therapies due to the limited insights into the mechanisms underlying tissue wasting. Here, we utilise a YAP-driven model of liver cancer in zebrafish, which rapidly develops cachexia, to uncover an evolutionarily conserved role for bile acid disruption in the onset of cachexia. Spatial transcriptomic analysis revealed that YAP induces a bi-lineage cholangiocarcinoma phenotype, which was associated with bile acid dysregulation. Mechanistically, we establish that both bile acid synthesis (via CYP7A1) and signalling through the bile acid receptor TGR5 are essential for cachexia induction. Notably, we find that the promotion of bile acid excretion with odevixibat ameliorates cachexia. Together, our findings reveal an evolutionarily conserved mechanism by which YAP promotes cachexia and suggest a potential therapeutic strategy to treat the syndrome.

## Introduction

Cachexia is a complex catabolic syndrome characterised by profound loss of adipose and muscle tissue^1,2^. The recognition of cachexia dates to ancient Greek times, where the Hippocratic Corpus described the wasting that defines this condition as “the flesh is consumed and becomes water”. Although cachexia can complicate a broad range of malignancies, it is particularly prevalent in patients with gastrointestinal cancers, including liver cancer^3^. Tumours contribute to this wasting phenotype by secreting circulating cachectic factors that promote the breakdown of fat and muscle stores, releasing fatty acids and amino acids that support tumour growth^1^. Despite extensive research, the molecular underpinnings of cancer-associated cachexia remain incompletely defined. Although often associated with late-stage disease, studies have shown that cachexia can precede the clinical detection of malignancies and is strongly associated with adverse survival outcomes^2–4^. Despite the current lack of effective therapies for cachexia, the promising results of ponsegromab (anti-GDF15) in recent phase 2 clinical trials represent a pivotal advance, that may herald a new era for the development of targeted treatments to combat cancer-associated cachexia^5^.

The Hippo signalling network is a conserved pathway that regulates growth control via the transcriptional regulator Yes-associated protein (YAP)^6,7^. YAP is a key oncogene in many solid tumours, including cholangiocarcinoma^6,8–10^. Our previous studies have revealed that YAP reprograms metabolism to stimulate *de novo* nucleotide biosynthesis and lipogenesis, which supports oncogenic growth^11–13^. It is now recognised in the field that YAP is a master regulator of metabolism that fuels tumorigenesis^14–16^. Recent studies have identified that YAP can drive tumourigenesis via non-cell-autonomous mechanisms, including cell competition^17,18^. Despite these insights, the impact that YAP may have in systemic metabolic conditions such as cancer-associated cachexia is poorly understood.

In this study, we uncover a previously unappreciated role for YAP in driving cancer-associated cachexia. Using zebrafish models, we show that YAP acts as a potent inducer of liver cancer-induced cachexia. Transcriptomic approaches reveal that YAP disrupts the FXR-regulated network of genes responsible for maintaining bile acid homeostasis. Mechanistically, we identify an essential role for the bile acid receptor TGR5 in the initiation of cachexia and show that reducing circulating bile acid levels with odevixibat alleviates cachexia in tumour-bearing fish. Finally, we provide evidence that YAP-driven bile acid signalling is sufficient to induce cachexia in mice. Together, these studies identify bile acid disruption and TGR5 signalling as key mediators of cancer-associated cachexia and reveal potential therapeutic opportunities for intervention.

## Results

### Development of a YAP-driven model of liver cancer that exhibits cachexia

To establish a model of liver cancer–associated cachexia, we generated a compound transgenic zebrafish line with hepatocyte-specific expression of a constitutively active form of YAP (Tg(*fabp10a*:yap1^S87A^;*cryaa*:Venus))^11^, on a loss-of-function mutant p53 background (*tp53*^M214K/M214K^)^19^, herein referred to as P53KO;*lf*:YAP (Fig. 1a). For all experiments, the lines were crossed onto a hepatocyte reporter Tg(*fabp10a*:CFP-NTR), hereafter referred to as *lf*:CFP, to visualise the liver. Initially, we observed that P53KO;*lf*:YAP fish showed a significant survival defect compared to wild-type (WT) controls, with approximately 40% loss in survival between 14- and 28-days post-fertilisation (dpf) (Fig. 1b). Notably, the P53KO fish and *lf*:YAP fish did not experience this level of mortality (Fig. S1a), indicating a synergistic effect between loss of p53 and overexpression of YAP. At 21 dpf, P53KO;*lf*:YAP fish displayed a bimodal phenotype, with one subgroup developing liver hyperplasia (hereafter referred to as PY-H), and another subgroup developing visible liver tumours (hereafter referred to as PY-TB) (Fig. 1c). PY-TB fish showed clear signs of wasting compared to WT fish and PY-H siblings, including contracted muscle width and a reduction in visceral adipose tissue, as seen with Nile Red staining (Fig. 1c). The wasting phenotype was not observed in P53KO or *lf*:YAP fish (Fig. S1b). Next, biometric analysis of large cohorts of WT and P53KO;*lf*:YAP fish was conducted with liver, muscle, and adipose tissue normalised to the standard length (SL) of each fish. Using this approach, we confirmed that PY-TB fish had significant liver enlargement (Fig. 1f), alongside a reduction in muscle (Fig. 1e) and adipose (Fig. 1g) tissue. Together, these studies suggest that hepatic YAP activation in the context of *tp53* loss is sufficient to drive systemic wasting consistent with cachexia.

**Figure 1.**
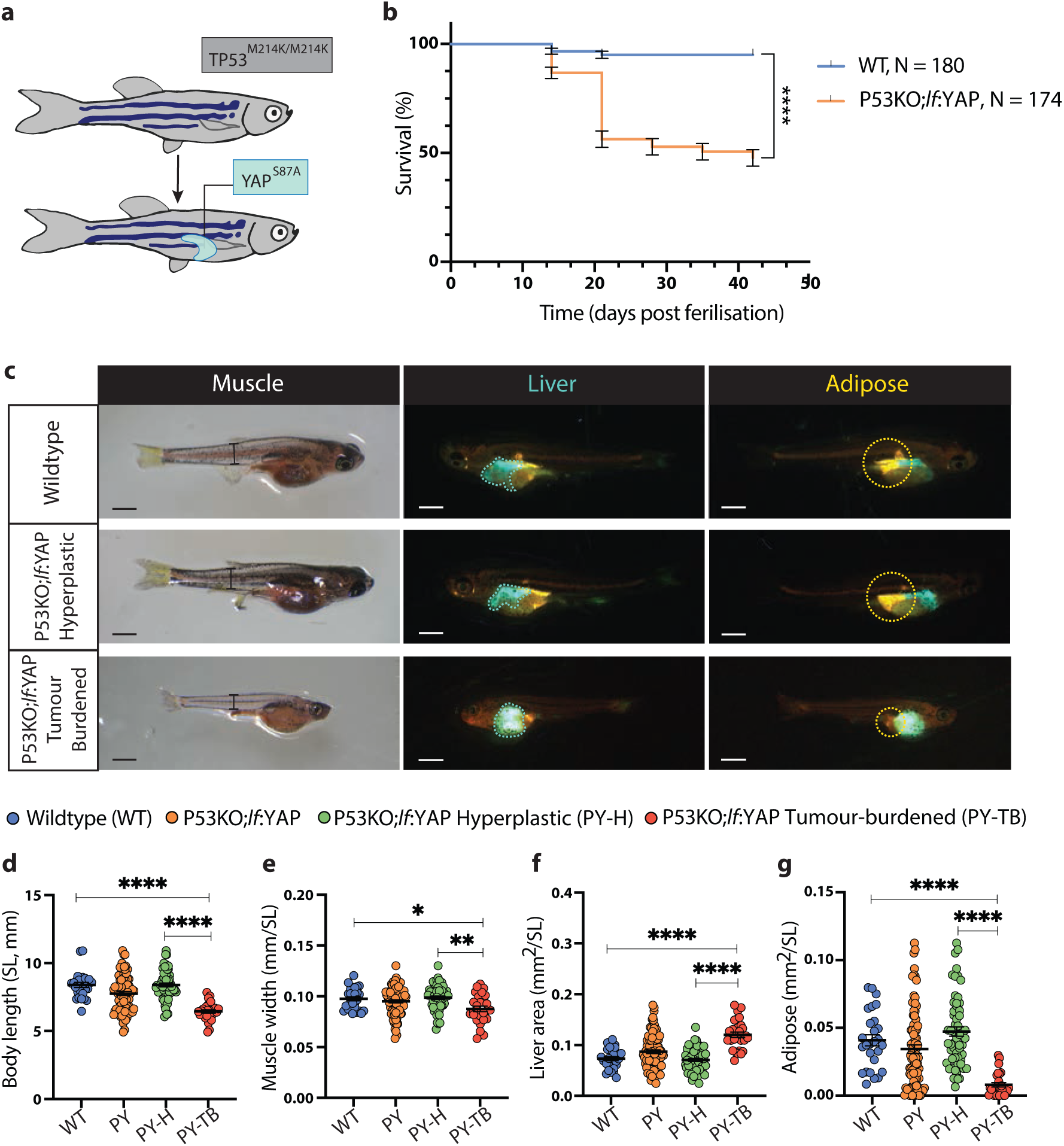
Development of a YAP-driven model of liver cancer that exhibits cachexia. **a**, Schematic of the double transgenic model, P53KO;*lf*:YAP (PY), with global knockout of the tumour suppressor gene tp53 and liver-specific expression of constitutively active YAP. **b**, Kaplan–Meier survival curves of wild-type (WT; n = 180) and PY zebrafish (n = 174). **c**, Representative images of 21 dpf WT and PY zebrafish, including hyperplastic (PY-H) and tumour-burdened (PY-TB) phenotypes, showing gross morphology (brightfield), lf:CFP-NTR liver fluorescence (cyan) and Nile Red-stained adipose tissue (yellow). In brightfield images, black lines indicate muscle width measurement. Scale bar, 1 mm. **d–g**, Biometrics of body length (**d**, standard length, SL), muscle width (**e**), liver size (**f**) and adipose tissue size (**g**) in WT, PY, PY-H and PY-TB zebrafish at 21 dpf. All data are mean ± s.e.m. P values were determined by log-rank (Mantel–Cox) test (b), one-way ANOVA with Tukey’s multiple-comparisons test (d,e) and Kruskal–Wallis test with Dunn’s multiple-comparisons test (f,g). *P < 0.05, **P < 0.01, ***P < 0.001, ****P < 0.0001.

### Spatial transcriptomic analysis reveals a signature of liver cancer-associated cachexia

To define the spatial architecture of liver cancer-associated cachexia, we performed spatial transcriptomic profiling using BGI Stereo-seq technology^20,21^. We collected cryosections from WT and PY-TB zebrafish at 21 dpf, aligning spatial gene expression with adjacent H&E-stained sister sections (Fig. 2a). Spatial transcriptomic spots were grouped into distinct domains based on similarities in gene expression profiles, using the Seurat pipeline with principal component analysis and graph-based clustering (Fig. 2b). Cluster identities were defined by the Seurat pipeline based on regionally enriched marker genes, with likely tissue or cell type interpretation informed by Daniocell (Fig. 2c, Fig. S2a–c)^22^. To focus specifically on hepatocyte clusters, we identified spatial spots with high expression of the hepatocyte marker *fabp10a* (Fig. 2d). Hepatocyte-enriched spots were assigned to their respective samples based on spatial coordinates, enabling stratification of WT (blue spots) or PY-TB (red spots). We then performed differential gene expression (DEG) analysis between WT and PY-TB hepatocyte clusters (Fig. 2e). Amongst the DEGs, PY-TB hepatocyte clusters showed significant downregulation of hepatocyte-specific markers (*hpxa*, *ahsg2*, *plg*) and upregulation of cholangiocyte-associated genes (*anxa4*, *lgals2b*, *cldn15lb*, *tm4sf4*), as well as canonical YAP targets (*amotl2a*, *amotl2b*) (Fig. 2e). Spatial mapping of *anxa4* expression confirmed broad elevation across PY-TB hepatocyte clusters, indicating ectopic activation of cholangiocyte-associated programmes within the tumour-burdened liver parenchyma (Fig. 2f). Gene set enrichment analysis (GSEA) confirmed enrichment of a canonical YAP signature amongst the PY-TB DEGs (Fig. 2g). Enrichment of a bile duct–associated cell identity signature, and a concomitant decrease in a hepatocyte identity signature (HNF4α signature) was also observed (Fig. 2h,i). To validate the transcriptional changes observed by Stereo-seq, bulk RNA-seq analysis was conducted on liver tissue isolated from WT and PY-TB zebrafish at 21 dpf. DEG analysis recapitulated core features of the spatial dataset, including upregulation of YAP targets, loss of hepatocyte markers and induction of cholangiocyte-associated markers (Fig. S2d). GSEA of the bulk RNA-seq data confirmed enrichment of both a YAP signature and a bile duct cell identity signature (Fig. S2e,f). Collectively, these studies suggest that YAP-transformed hepatocytes undergo a cell fate shift towards the cholangiocyte lineage, consistent with previous studies in mice^8^. Histological analysis revealed that PY-TB fish livers harboured cytological abnormalities (e.g., hepatocytes with increased nuclear to cytoplasm ratio), as well as architectural disruptions (e.g., glandular morphology), reminiscent of a combined hepatocellular carcinoma-cholangiocarcinoma phenotype (Fig. 2j). To validate the YAP-driven changes in liver cell fate we performed immunofluorescence and confirmed increased expression of the cholangiocyte marker ANXA4 in PY-TB livers (Fig. 2j). Importantly, we observed bi-lineage cells with co-expression of hepatocyte (GFP) and cholangiocyte (ANXA4) markers, suggesting that YAP induces lineage infidelity and transdifferentiation of hepatocytes to cholangiocytes^8^. In contrast, histological sections derived from PY-H fish exhibited minimal cystic distortion and no sign of bi-lineage ANXA4+ hepatocytes (Fig. S2f). Collectively, these findings suggest that YAP-driven tumour initiation is associated with lineage infidelity as transformed hepatocytes acquire cholangiocyte identity.

**Figure 2.**
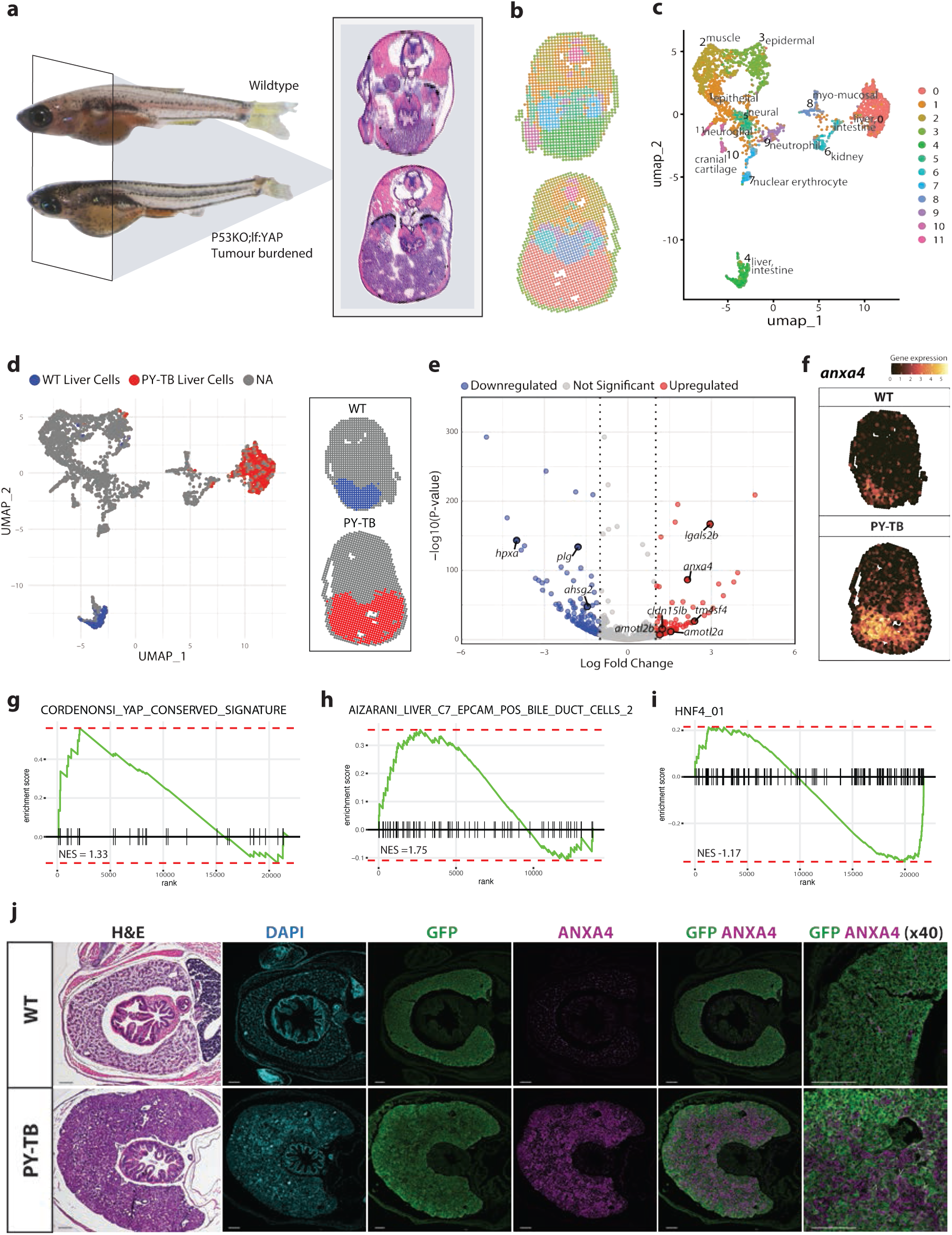
Spatial transcriptomic analysis reveals a signature of liver cancer-associated cachexia. **a**, Schematic of the spatial transcriptomics (SRT) workflow. Cryosections were obtained from the indicated anatomical region (highlighted in schematic). Adjacent sister sections were stained with H&E, and sections for SRT were processed with BGI Stereo-seq technology. **b**, Spatial feature plot showing spot-level transcriptomic clustering of WT and PY-TB zebrafish samples. Clusters were identified by Seurat using principal component analysis and graph-based clustering of transcriptomic neighbourhoods. **c**, Integrated UMAP of all spatial transcriptomic spots coloured by Seurat-defined cluster identity. Cluster annotation was guided by regionally enriched marker genes and tissue interpretation using Zebrahub. **d**, UMAP and spatial feature plots highlighting hepatocyte-enriched spots defined by expression of hepatocyte marker, *fabp10a*, above the 75th percentile. Cells are grouped by sample: WT hepatocytes (blue) and PY-TB hepatocytes (red). **e**, Volcano plot of differentially expressed genes (DEGs) between PY-TB hepatocytes and WT hepatocytes, both defined by *fabp10a* expression in the spatial data. **f**, Spatial projection of cholangiocyte marker, *anxa4,* expression, visualised over spatial coordinates of the tissue section. **g-i**, Gene set enrichment analysis (GSEA) plots of selected pathways enriched in PY-TB hepatocytes versus WT, derived from DEGs identified in the SRT dataset. **j**, H&E and immunofluorescence staining of liver sections of WT and PY-TB fish at 21 dpf. Nuclei are marked with DAPI (cyan), hepatocytes with GFP (green), and cholangiocytes with ANXA4 (magenta). White arrow represents GFP+/ANXA4+ bi-lineage cells. Scale bar, 100 μm.

### YAP disrupts bile acid homeostasis and drives muscle and adipose wasting

We next sought to assess the systemic consequences of tumour burden. To this end, high-resolution 3D micro-CT was performed on WT, PY-H and PY-TB samples at 21 dpf. PY-TB fish displayed hepatomegaly alongside substantial reductions in both adipose and skeletal muscle tissue relative to WT and PY-H counterparts (Fig. 3a). Quantitative volumetric assessment of CT scans derived from an extended cohort confirmed these findings, revealing significantly increased liver volume (Fig. 3b), decreased muscle cross-sectional area (Fig. 3c), and severe adipose depletion (Fig. 3d). To examine the visceral adipose depot at cellular resolution, we performed confocal imaging of Nile Red-stained adipose tissue. We found that PY-TB fish harboured fewer adipocytes and a shift in the size distribution towards smaller adipocytes (Fig. 3e,f, Fig. S3a). To further investigate changes in body composition, we performed a BrdU incorporation assay to assess tissue growth dynamics. Using this method, we observed increased BrdU incorporation in the liver of both PY-H and PY-TB fish compared to WT, consistent with YAP activation stimulating tissue growth. (Fig. S3b, S3c). Interestingly, a significant decrease in BrdU incorporation in the muscle of PY-TB compared to PY-H and WT fish was observed (Fig. S3d, S3e). We assessed mTORC1 activation and found a reduction in phospho-S6 (pS6) staining in muscle tissue of PY-TB compared to WT fish, indicating reduced anabolic mTORC1 activity (Fig S3f). To determine whether these muscle phenotypes were accompanied by transcriptional reprogramming, we interrogated the myogenic cluster in our spatial transcriptomics dataset. Cluster identity was validated using marker gene expression and anatomical alignment with H&E-stained sections (Fig S3g). Compared to WT muscle, PY-TB muscle showed widespread downregulation of contractile and structural genes (*myhz1.3*, *myhc4*, *desma*) and suppression of lipid regulators (*lpin1*), along with increased expression of thermogenic (*ucp1*), metabolic stress (*igfbp1a*), and inflammatory (*c3a.3*) markers (Fig S3h). Spatial feature plots indicated decreased *myhz1.3* expression in PY-TB muscle relative to WT muscle, suggesting compromised muscle fibre integrity. Overall, these results reveal that YAP-driven tumour development is linked to significant changes in body composition, including reductions in muscle and adipose tissue - features typical of cancer-associated cachexia.

**Figure 3.**
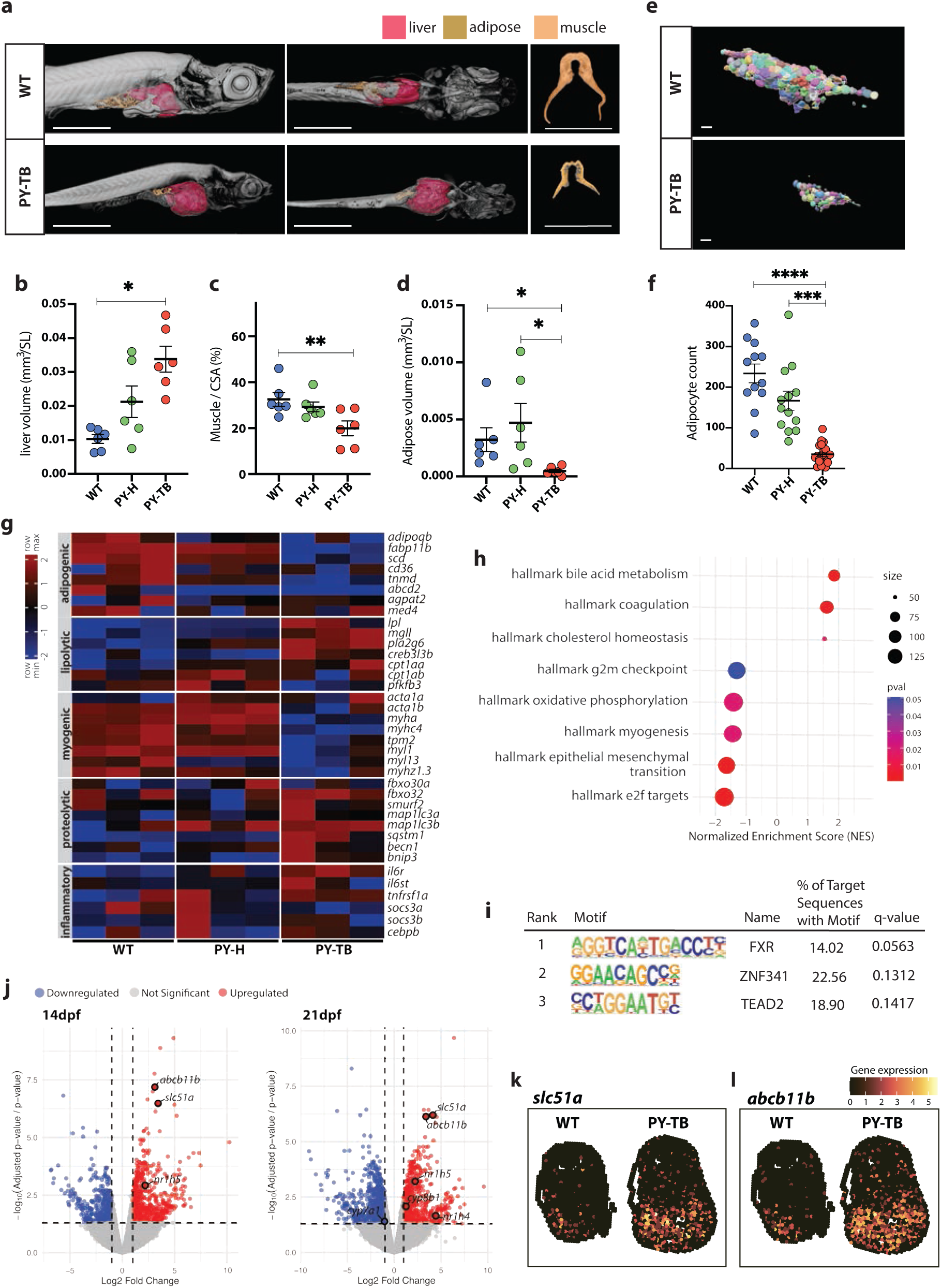
YAP disrupts bile acid homeostasis and drives muscle and adipose wasting. **a**, Representative 3D reconstructions and coronal sections from micro-CT scans of 21 dpf WT and PY-TB zebrafish. Liver, adipose and muscle tissues were segmented using Avizo3DPro software following image acquisition and volume reconstruction with phoenix datos|x. Liver, red; adipose, yellow; muscle, orange. Scale bar, 1 mm. **b**, Quantification of micro-CT-derived phenotypes in WT, PY-H and PY-TB zebrafish at 21 dpf: liver volume (**b**), muscle cross-sectional area normalized to total body cross-sectional area at the same slice (measured at the level of vertebra C8) (**c**), and adipose volume (**d**). **e**, Representative IMARIS-rendered reconstructions of confocal z-stacks showing Nile Red-stained adipose tissue in WT and PY-TB zebrafish at 21 dpf. Scale bar, 100μm. **f**, Quantification of adipocyte number from IMARIS analysis of confocal z-stacks of WT, PY-H and PY-TB zebrafish at 21 dpf. **g**, Heat map of selected adipogenic, lipolytic, myogenic, proteolytic and inflammatory gene expression profiles in pooled WT, PY-H and PY-TB zebrafish at 21 dpf, determined by bulk RNA-seq (n = 4 pools of 10 fish at 14 dpf; n = 3 pools of 10 fish at 21 dpf). **h**, Dot plot of significantly enriched HALLMARK pathways identified by GSEA of bulk RNA-seq comparing PY-TB to WT zebrafish at 21 dpf. **i,** The top three enriched transcription factor-binding motifs identified using Hypergeometric Optimization of Motif EnRichment (HOMER) motif analysis, among significantly upregulated genes upregulated in PY-TB versus WT zebrafish at 21 dpf, from bulk RNA-seq analysis. **j,** Volcano plot of DEGs from bulk RNA-seq, comparing PY-TB and WT zebrafish at 14 and 21 dpf. Bile acid-related genes are labelled. **k-l,** Spatial feature plots of bile acid-related transporter expression in WT and PY-TB tissue sections from SRT: *slc51a* (**k**; bile acid transporter) and *abcb11b* (**l**; bile salt export pump). All data are mean ± s.e.m. P values were determined by Kruskal–Wallis test with Dunn’s multiple-comparisons post hoc test (**b–d**), one-way ANOVA with Brown–Forsythe correction (**f**). *P < 0.05, **P < 0.01, ***P < 0.001, ****P < 0.0001.

To capture the global transcriptional landscape, bulk RNA-seq analysis was performed on pooled fish at 21 dpf. When compared to WT and PY-H fish, PY-TB fish exhibited a transcriptional profile characteristic of cachexia, with coordinated downregulation of adipogenic and myogenic genes, while lipolytic, proteolytic, and inflammatory genes were upregulated (Fig. 3g). GSEA of hallmark signatures derived from bulk RNA-seq data revealed a strong upregulation of genes associated with bile acid metabolism in PY-TB fish compared to WT (Fig. 3h), implicating bile acid dysregulation as a feature associated with cachexia. To identify the top-ranked transcription factors driving the cachectic response, we performed HOMER motif enrichment analysis on upregulated genes in PY-TB versus WT. Interestingly, the most enriched motif corresponded to farnesoid X receptor (FXR), a nuclear receptor activated by bile acids that governs the transcriptional regulation of bile acid homeostasis (Fig. 3i). Consistent with this, DEG analysis between WT and PY-TB fish showed perturbed expression of FXR-regulated genes involved in bile acid synthesis (*cyp7a1*, *cyp8b1*, *cyp27a1.2*) and bile acid transport (*slc10a2*, *abcb11b*, *slc51a*, *fabp6*) (Fig. 3j). These changes were apparent at both early (14 dpf) and late (21 dpf) stages, indicating a progressive disruption of bile acid homeostasis during tumour and cachexia development. Re-examining our spatial transcriptomic dataset, we found increased expression of *slc51a* and *abcb11b* in PY-TB livers compared to WT (Fig. 3k,l), consistent with FXR acting as a positive regulator of these bile acid transporters in zebrafish^23^. Together, these findings suggest tumour-driven bile acid dysregulation as a feature associated with the onset of cachexia in a YAP-driven tumour model.

### YAP-induced cachexia requires CYP7A1 and TGR5

To investigate whether bile acid dysregulation contributes to YAP-driven cachexia, we employed a CRISPR–Cas9 strategy that enabled multiplexed gene knockouts in F0 zebrafish. This method involved co-targeting the pigmentation gene *slc45a2*, which served as a marker for injection and editing efficiency, since loss of *slc45a2* results in an albino phenotype (Fig. S4a). Using this approach, we injected WT and P53KO;lf:YAP embryos at the one-cell stage with Cas9 complexed with synthetic CRISPR RNAs (crRNA) targeting exonic regions of *tgr5* or *cyp7a1*. TGR5 is a bile acid–responsive receptor expressed in peripheral tissues, while CYP7A1 is a rate-limiting enzyme in bile acid synthesis. We first confirmed successful indel efficiency by performing Sanger sequencing and Inference of CRISPR Editing (ICE) analysis on exon regions of *slc45a2*, *tgr5*, and *cyp7a1* within individual embryos (Fig S4b-g). In PY-TB fish, loss of *tgr5* (PY-TB;cr*slc45a2;*cr*tgr5*) significantly rescued key cachectic features observed in the control crispants (PY-TB;cr*slc45a2*) (Fig. 4a). For example, PY-TB;cr*slc45a2*;cr*tgr5* fish showed improved adipose retention at 21 dpf, without any observable reduction in tumour size (Fig. 4a). Similarly, loss of *cyp7a1* (PY-TB;cr*slc45a2*;cr*cyp7a1*) increased the size of the visceral adipose depot in the tumour-burdened group (Fig. 4a). Biometric quantification of large cohorts revealed significant increases in both muscle width and adipose area in PY-TB;cr*slc45a2*;cr*tgr5* and PY-TB;cr*slc45a2*;cr*cyp7a1* fish relative to PY-TB;cr*slc45a2* controls (Fig. 4b-e). Collectively, these studies provide genetic evidence that bile acid synthesis and activation of TGR5 receptors play a functional role in the induction of cancer-associated cachexia.

**Figure 4.**
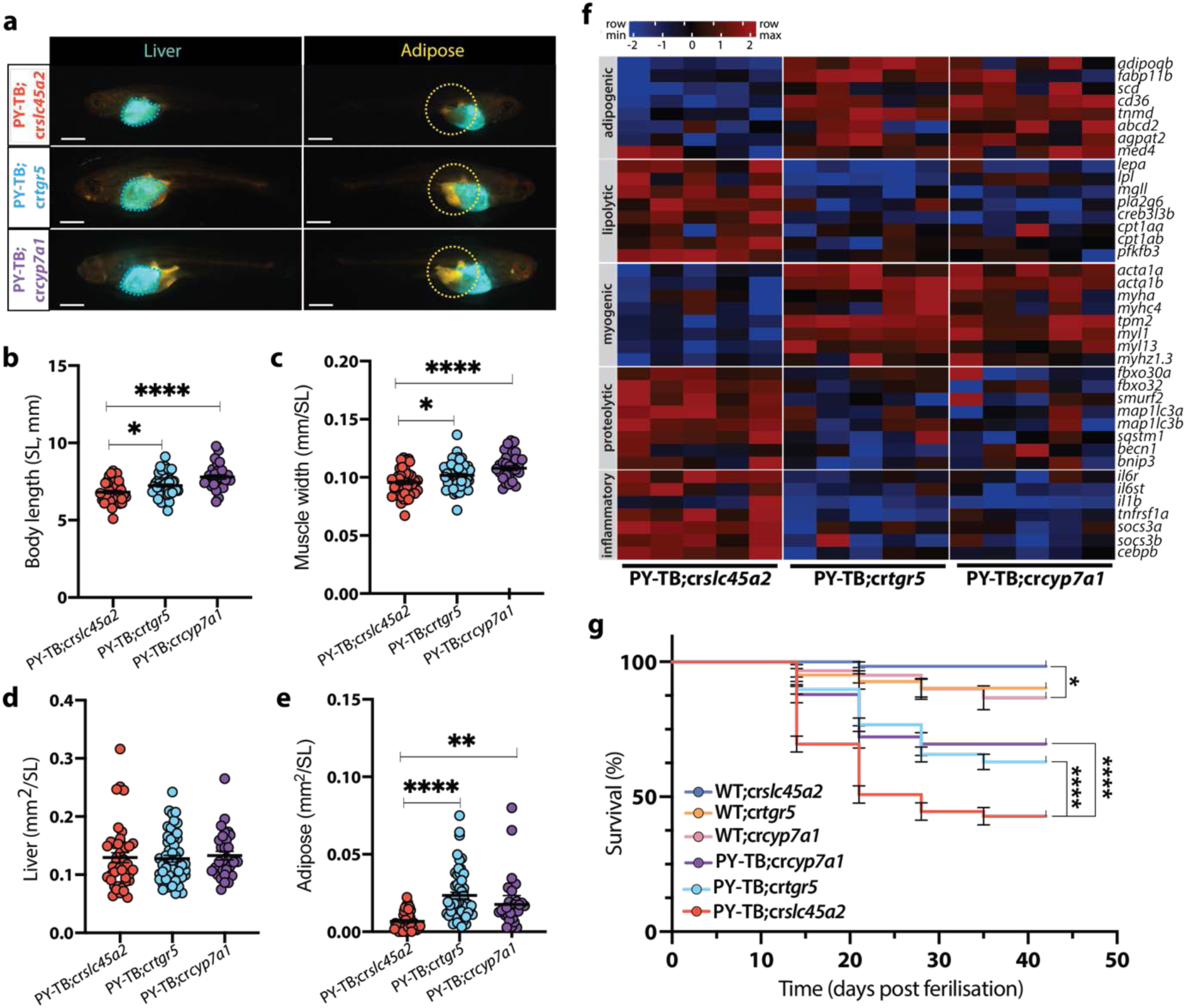
YAP-induced cachexia requires CYP7A1 and TGR5. **a**, Representative images of PY-TB;cr*slc45a2*), PY-TB;cr*tgr5* and PY-TB;*crcyp7a1* zebrafish at 21 dpf. Liver tissue, *lf*:CFP-NTR fluorescence (cyan); adipose tissue, Nile Red staining (yellow). Scale bar, 1 mm. **b-e**, Biometrics of body length (**b**, standard length, SL), muscle width (**c**), liver size (**d**) and adipose tissue size (**e**) in PY-TB;cr*slc45a2*, PY-TB;cr*tgr5* and PY-TB;cr*cyp7a1* zebrafish at 21 dpf. **f**, Heat map of selected adipogenic, lipolytic, myogenic, proteolytic and inflammatory gene expression profiles in PY-TB;cr*slc45a2*, PY-TB;cr*tgr5* and PY-TB;cr*cyp7a1* zebrafish at 21 dpf, determined by bulk RNA-seq (n = 5 pooled samples of 5 fish per condition). **g**, Kaplan–Meier survival curves of WT;cr*slc45a2* (n = 60), WT;cr*tgr5* (n = 82), WT;cr*cyp7a1* (n = 60), PY-TB;cr*slc45a2* (n = 236), PY-TB;cr*tgr5* (n = 283) and PY-TB;cr*cyp7a1* (n = 115) zebrafish. All data are mean ± s.e.m. P values were determined by one-way ANOVA with Tukey’s multiple-comparisons test (**b,c**), Kruskal–Wallis test with Dunn’s multiple-comparisons post hoc adjustment (**d,e**), and log-rank (Mantel–Cox) test for survival analysis (**g**). *P < 0.05, **P < 0.01, ***P < 0.001, ****P < 0.0001.

To determine whether these phenotypic effects reflect underlying transcriptional changes, we performed bulk RNA-seq on pooled PY-TB samples for each genotype. Remarkably, loss of either *tgr5* or *cyp7a1* caused a significant reduction in the canonical cachectic gene signature (Fig 4f). Specifically, we observed increased expression of adipogenic and myogenic genes, and decreased expression of inflammatory, lipolytic, and proteolytic genes in PY-TB;cr*slc45a2*;cr*tgr5* and PY-TB;cr*slc45a2*;cr*cyp7a1* fish compared to control PY-TB;cr*slc45a2* fish (Fig. 4f). Notably, these molecular and phenotypic rescues translated into better survival.

Kaplan–Meier analysis revealed a significant improvement in survival outcomes in PY-TB fish lacking either *tgr5* or *cyp7a1*, compared to PY-TB fish (Fig. 4g). To check if these effects were specific to the tumour context, we examined the impact of *tgr5* or *cyp7a1* loss in a WT background. In WT fish, loss of *tgr5* slightly increased muscle but had no significant impact on other measures (Fig. S4h-k). Conversely, loss of *cyp7a1* in WT fish reduced adiposity, consistent with impaired bile acid–mediated lipid absorption (Fig. S4h-k). Transcriptional analysis of WT crispants showed modest upregulation of adipogenic genes in both backgrounds, whereas *cyp7a1* crispants showed a subtle upregulation of myogenic and proteolytic genes (Fig. S4L). Overall, these findings demonstrate that YAP-driven cachexia requires both bile acid synthesis and TGR5-mediated signalling. The coordinated recovery of muscle and fat wasting, gene expression profiles, and improved survival upon loss of either *tgr5* or *cyp7a1* support a model where YAP-driven tumourigenesis reprograms systemic metabolism via bile acid–TGR5 signalling.

### Odevixibat restores bile acid homeostasis and prevents cancer-associated cachexia

Having shown that bile acid signalling is necessary to induce cachexia in a YAP-driven model of liver cancer, we next sought to determine whether bile acids alone are sufficient to cause wasting. To evaluate the systemic effects of excess bile acids, WT zebrafish were treated with taurocholic acid (TCA) from 18–25 dpf (Fig. S5a). TCA treatment led to a significant reduction in adipose tissue compared to DMSO controls (Fig. S5b-f), indicating a lipolytic effect driven by bile acid overload. Odevixibat is an ileal bile acid transporter (SLC10A2) inhibitor that blocks bile acid reabsorption and has recently been approved for treating progressive familial intrahepatic cholestasis^24,25^. To determine whether pharmacological inhibition of bile acid reabsorption could mitigate the detrimental effects of excess bile, fish were treated with odevixibat (Odev), either alone or combined with TCA. Notably, co-treatment significantly prevented TCA-induced adipose loss, restoring adipose levels to those observed in untreated WT fish, whereas odevixibat alone had no significant impact on adipose tissue. Transcriptomic profiling of pooled samples revealed that TCA treatment led to an enrichment of the hallmark bile acid metabolism signature (Fig. S5g). In contrast, co-treatment with TCA and odevixibat reversed this transcriptional pattern, suppressing the hallmark bile acid metabolism signature and enhancing the hallmark adipogenesis signature when compared to TCA-treated counterparts (Fig. S5h). Together, these findings indicate that bile acid accumulation is sufficient to induce adipose wasting, and that odevixibat treatment can mitigate the effects of bile acid overload.

To determine whether pharmacological suppression of bile acid reabsorption could inhibit cancer-associated cachexia, PY-TB fish were treated with odevixibat from 14–21 dpf, a period corresponding to early tumour development and the onset of systemic wasting (Fig. 5a). Compared to DMSO-treated PY-TB controls, odevixibat-treated PY-TB fish showed a remarkable rescue of systemic wasting, with notable improvements in muscle width and adipose tissue (Fig. 5b-f). Bulk RNA-seq analysis of PY-TB fish treated with odevixibat revealed a suppression of the cachectic signature relative to PY-TB controls (Fig. 5g). Specifically, we observed an increase in adipogenic and myogenic genes, along with a decrease in the expression of lipolytic, proteolytic, and inflammatory genes (Fig. 5g). GSEA confirmed a repression of the hallmark bile acid metabolism signature in odevixibat-treated fish (Fig. S5I), supporting a model in which odevixibat inhibits bile acid-driven cachexia. Finally, Kaplan–Meier analysis across the treatment period showed significantly improved survival in odevixibat-treated PY-TB fish cohorts (Fig. 5h). Collectively, these findings demonstrate that odevixibat can reduce bile acid overload and slow the progression of cancer-associated cachexia.

**Figure 5.**
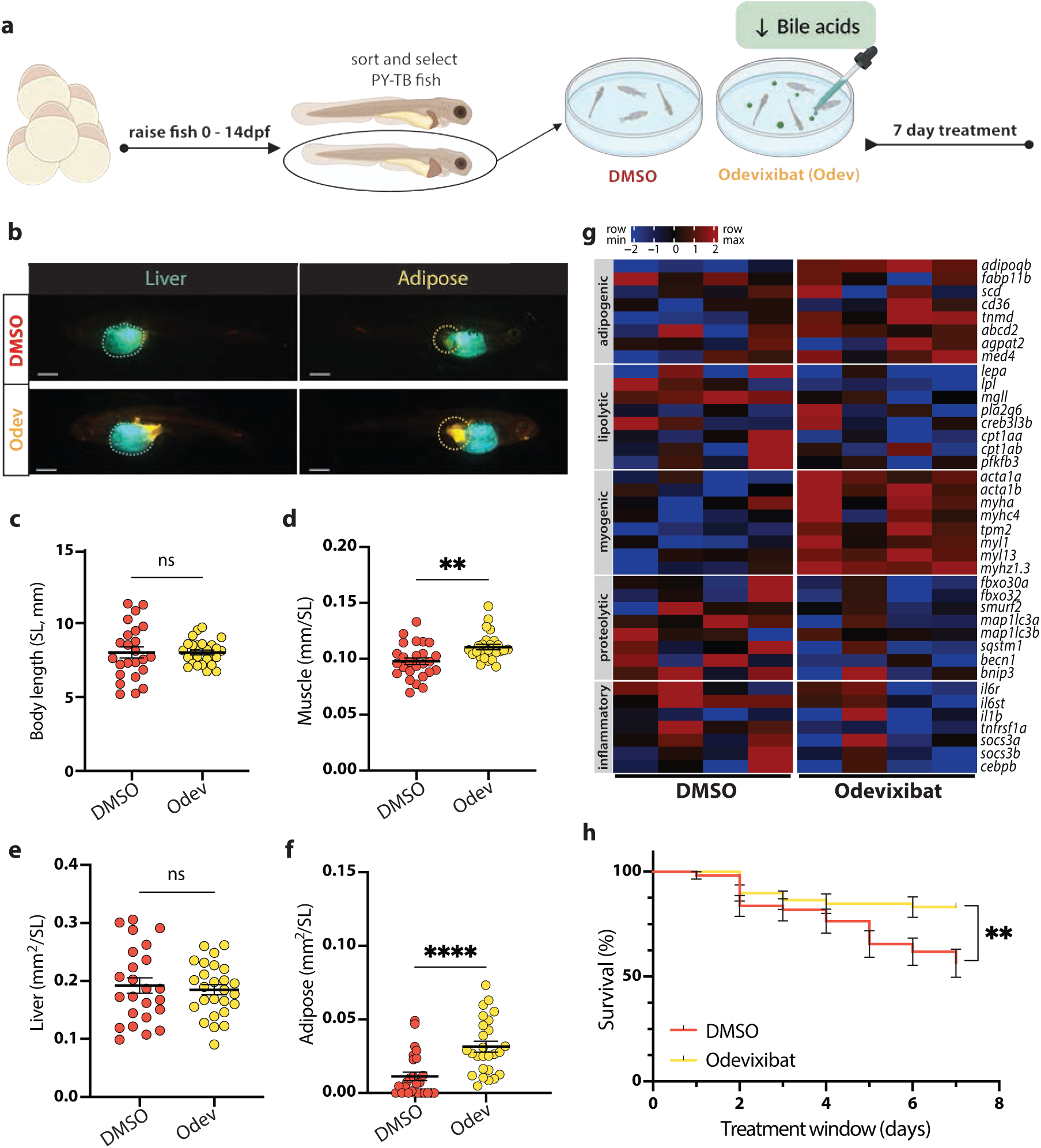
Odevixibat restores bile acid homeostasis and prevents cancer-associated cachexia. **a**, Schematic of the Odevixibat (Odev) treatment intervention protocol. **b**, Representative images of control (DMSO) and Odev-treated 21 dpf PY-TB zebrafish. Liver tissue, *lf*:CFP-NTR fluorescence (cyan); adipose tissue, Nile Red staining (yellow). Scale bar, 1 mm. **c-f**, Biometrics of body length (**c**, standard length, SL), muscle width (**d**), liver size (**e**) and adipose tissue size (**f**) in 21 dpf PY-TB DMSO and Odev-treated zebrafish. **g**, Heat map of selected adipogenic, lipolytic, myogenic, proteolytic and inflammatory gene expression profiles in pooled PY-TB DMSO and Odev-treated zebrafish at 21 dpf, determined by bulk RNA-seq (n = 4 pooled samples of 5 fish per condition). **h**, Kaplan–Meier survival curves of PY-TB DMSO (n = 55) and Odev-treated (n = 59) zebrafish over the 7-day treatment window. All data are mean ± s.e.m. P values were determined by Welch’s t-test (**c**), Mann–Whitney test (**d,f**), unpaired two-tailed t-test (**e**) and log-rank (Mantel–Cox) test for survival analysis (**g**). *P < 0.05, **P < 0.01, ***P < 0.001, ****P < 0.0001.

### YAP-induced cachexia is evolutionarily conserved

Having shown that YAP potently induces cancer-associated cachexia in zebrafish, we were motivated to investigate whether this mechanism was conserved in mice. To achieve this, a hepatocyte-specific doxycycline (Dox) inducible YAP overexpression model was utilised to monitor the body mass of mice over time (Fig 6a)^8^. Examination of the body weight over time revealed that YAP induction for 2 weeks resulted in a 15% reduction in body mass (Fig 6b). Interestingly, this mouse model was recently shown to develop hepatomegaly and cholestasis, with a significant increase in serum bile acids^26^. To determine whether disruption of bile acid metabolism was necessary to induce cachexia in mice, hepatocyte-specific Adeno-Associated Viral (AAV) vectors were deployed to overexpress YAP in the presence or absence of the bile acid exporter BSEP, which is encoded by the *Abcb11* gene (Fig 6c). Using this method, we found that YAP caused a 12% decrease in body mass that was restored by the concurrent overexpression of BSEP (Fig 6d). To confirm that lowering bile acid levels was sufficient to rescue cachexia, mice with hepatocyte-specific YAP overexpression were fed chow containing 2% cholestyramine resin, which enhances bile acid excretion (Fig 6c)^26^. Remarkably, the reduction in body mass observed following YAP overexpression was also rescued by cholestyramine resin (Fig 6e). In conclusion, these data demonstrate that YAP induces cachexia in a manner that is conserved across evolution.

**Figure 6.**
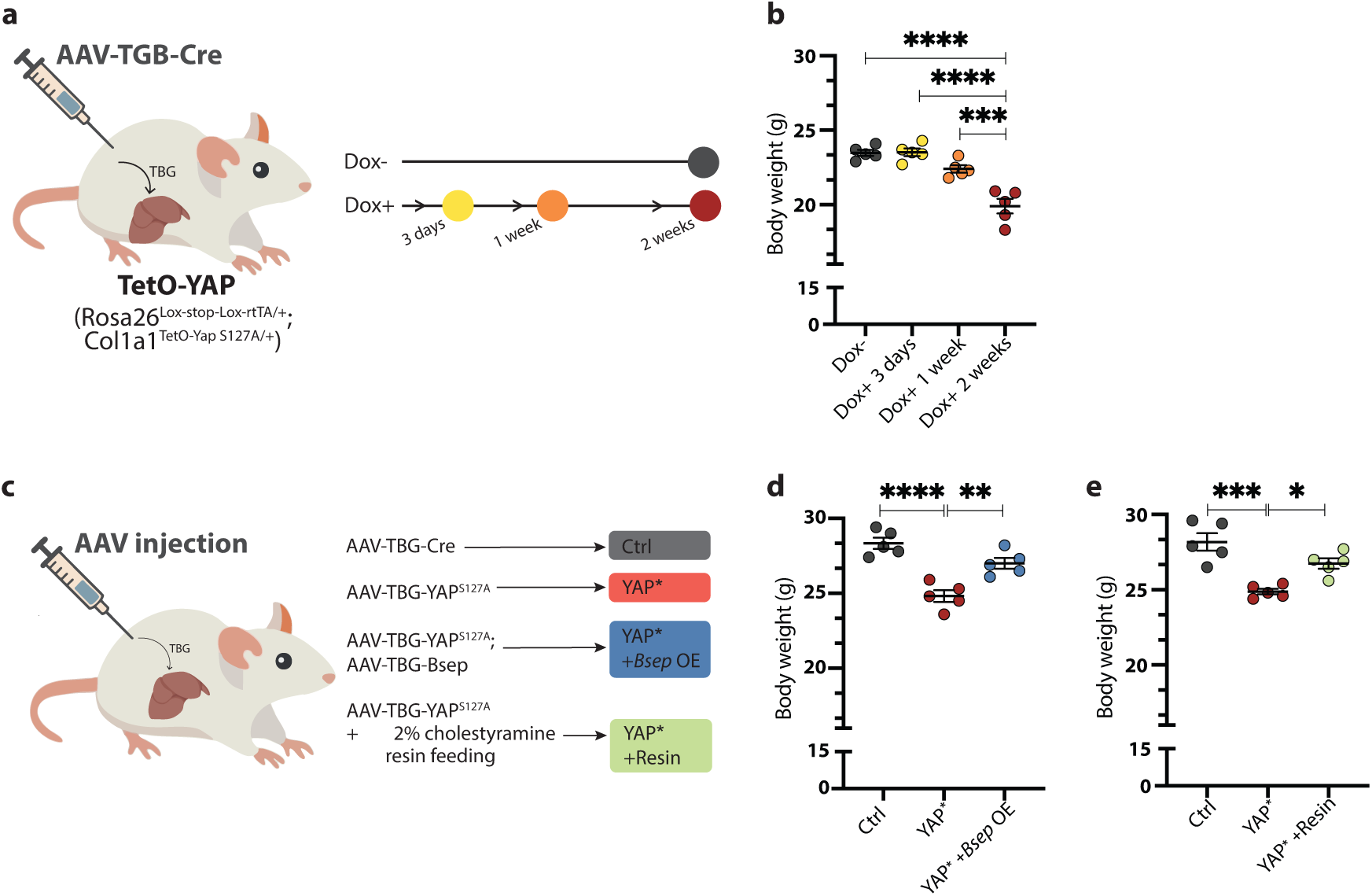
YAP-induced cachexia is evolutionarily conserved. **a**, Schematic of the experimental procedure enabling hepatocyte-specific Dox-inducible YAP activation (*AAV-TBG-Cre* injection into *Rosa26^Lox-stop-Lox-rtTA^;Col1a1^TetO-YAPS127A^*mice). **b**, Body mass determination in *AAV-TBG-Cre* injected *Rosa26^Lox-stop-Lox-rtTA^;Col1a1^TetO-YAPS127A^*mice over 2 weeks following Dox exposure. **c**, Schematic of the experimental procedure outlining the approach to genetically (*AAV-TBG-Bsep*) or therapeutically (2% cholestyramine resin) restore bile acid homeostasis. **d**, Body mass determination in WT, *AAV-TBG-YAP^S127A^* and *AAV-TBG-YAP^S127A^;AAV-TBG-Bsep* mice after 8 weeks. **e**, Body mass determination in WT, *AAV-TBG-YAP^S127A^*and *AAV-TBG-YAP^S127A^* fed 2% cholestyramine resin after 8 weeks. All data are mean ± s.e.m. P values were determined by Welch’s t-test. **P < 0.01, ***P < 0.001, ****P < 0.0001.

## Discussion

In summary, we have identified that YAP triggers cachexia via disruption of bile acid homeostasis. Mechanistically, we provide evidence that YAP-induced cachexia requires TGR5 signalling and can be suppressed by odevixibat, a recently approved SLC10A2 inhibitor that enhances bile acid excretion^24,25^. Collectively, these findings define cachexia as a conserved feature of YAP-driven tumours that is amenable to therapeutic intervention.

Our body of work builds on emerging evidence that YAP regulates cancer-associated cachexia. Previous studies in Drosophila have shown that Yorkie (the fly homologue of YAP) can induce a cancer cachexia-like wasting syndrome that affects muscle, adipose, and gonadal tissues^27,28^. In these models, the circulating factor ImpL2 (fly homologue of IGFBP) was found to be both necessary and sufficient to induce wasting by suppressing insulin signalling. Recent insights into Epithelioid hemangioendothelioma (EHE), a rare sarcoma caused by a fusion of YAP paralogue TAZ, WWTR(TAZ)-CAMTA1, revealed that plasma levels of the cachectic factor GDF-15 correlated with disease aggressiveness^29^. Given these studies, it is tempting to speculate that cachexia may be a hallmark of YAP-driven malignancies.

The Hippo-YAP pathway is now recognised as one of the key signalling cascades that drives the formation of cholangiocarcinoma^8,9,30–32^. In this context, YAP activation is rarely driven by mutations in the Hippo pathway^33^, but rather by environmental cues, including nutrient availability and mechanotransduction, both of which are dysregulated in chronic liver disease (i.e., MASLD, fibrosis, and cirrhosis^10,34^). At diagnosis, patients with cholangiocarcinoma can present with pruritus and unintentional weight loss, clinical features associated with elevated bile acids and cachexia, respectively. Indeed, the levels of bile acids, such as tauroursodeoxycholic acid (TUDCA), are increased in the serum of patients with cholangiocarcinoma^35^. Recently, Bardeesy and colleagues have identified distinct subtypes of cholangiocarcinoma, each with unique attributes^36^. Among these subtypes, a “Bi-lineage R3” group was characterised by increased dependence on HNF1B, bi-lineage features consistent with cells exhibiting both hepatocyte and cholangiocyte fates, and a gene expression profile enriched for bile acid secretion. Taken together, these studies suggest that our YAP-driven zebrafish model recapitulates many of the features of the Bi-lineage R3 subtype of cholangiocarcinoma.

Bile acid dysregulation has been associated with the pathophysiology of disease since ancient Greek times, as described in Hippocrates’ humoral theory of disease. Bile acids are now recognised to play a key role in regulating metabolism and energy expenditure via the receptors FXR and TGR5^37–41^. While FXR-dependent regulation of bile acid metabolism is highly conserved across vertebrates, evolutionary distinctions exist between species. For example, FXR activation enhances BSEP (*abcb11b*) expression in zebrafish but suppresses BSEP/*ABCB11* in the mammalian liver^38–40^. However, our findings demonstrate a close link between YAP and bile acid metabolism, consistent with previous studies^26,42–44^. For example, there is strong evidence that bile acids are sufficient to activate YAP in both hepatocytes and cholangiocytes^42,43^. Yet, it is also clear that YAP hyperactivation in the liver is sufficient to elevate bile acid levels^26,44^. The emerging consensus in the field suggests that during normal homeostasis, YAP activity is low due to FGFR-driven stimulation of the Hippo pathway, whereas upon exposure to environmental cues that inhibit the Hippo pathway, YAP alters FXR activity to stimulate bile acid production. In the context of tissue injury and regeneration, the interplay of YAP and bile acids represents an adaptive mechanism to re-establish tissue homeostasis (the so-called hepatostat model)^45,46^. However, in the context of tumourigenesis, when YAP is constitutively hyperactive, the disruption of bile acid homeostasis influences FXR and TGR5 activity, driving both tumour formation and the induction of cachexia.

The central dogma of cancer involves a framework wherein cells acquire oncogenic hallmarks that are primarily cell autonomous or focused on the tumour microenvironment^47^. Recently, there has been a growing recognition that addressing the systemic factors in cancer will provide a deeper understanding of the pathogenesis and may lead to new therapeutic approaches^2^. Consequently, defining the mechanisms underpinning cancer cachexia may identify new biomarkers and therapies to preserve metabolic homeostasis. Our studies provide strong evidence that circulating bile acids may play a key role in the interorgan communication that drives cachexia. Given that circulating bile acids already show potential clinical utility as biomarkers in liver diseases such as cholangiocarcinoma^35,40,48^, it will be of interest in future studies to ascertain if specific bile acid species play a dominant role in triggering cachexia. Moreover, we identified that the SLC10A2 inhibitor odevixibat is an effective therapy to suppress YAP-driven liver cancer cachexia. With this in mind, it will be interesting to explore whether odevixibat has efficacy in other models of cancer-associated cachexia. Finally, if the link between YAP and cachexia is common, then it may warrant investigation of recently developed TEAD inhibitors^49,50^, as putative anti-cachectic therapies.

## Materials and Methods

### Zebrafish husbandry

Zebrafish were maintained according to institutional Animal Experimentation Ethics Committee (AEEC) guidelines (Approval E634, E666, AEC2024-03). The following zebrafish lines were used in this study, as previously described: Tg(*-2.8fabp10a*:yap1^S87A^;*cryaa*:Venus)^11^, hereafter referred to as *lf*:YAP; Tg(*fabp10a*: CFP-NTR)^51^, hereafter referred to as *lf*:CFP; *tp53^M214K/M214K^* ^19^, hereafter referred to as P53KO. Compound transgenics were generated by crossing the aforementioned lines to achieve homozygosity. For all experiments, clutch siblings were used as controls, and all embryos and larvae were maintained at 28 °C. All zebrafish experiments were performed at the larval stage, and therefore, the sex of the organism was not yet determined.

### Mice husbandry

Mouse protocols and procedures were approved by the Harvard Medical School Institutional Animal Care and Use Committee (HMS-IACUC). Mice were housed in a pathogen-free facility under a 12 h light/dark cycle. All mice were fed ad libitum with LabDiet® 5053. In the experiments, the maximum tumour burden and weight loss were approved by HMS-IACUC, and the burden did not exceed the standard limits. The timeline of YAP induction and phenotyping of the liver has been described previously^26,32^. In this study, the endpoints selected for the current mouse models reflect the minimum duration needed to observe moderate liver enlargement and tumour formation. No mice exhibited severe visible illness that met the veterinarian’s criteria for euthanasia at the endpoint. All mice were sacrificed between Zeitgeber Time (ZT) 5–8. Both male and female mice were used in the experiments. *Rosa26^lox-stop-lox-rtTA/+^*; *Col1a1^Teto^*^-YapS127A/+^ (TetO-YAP) mice were obtained from Dr. Fernando Camargo’s lab^8^.

### Survival Analysis

To generate survival curves, zebrafish clutches were collected and stored in petri dishes containing E3 water (5 mM NaCl, 0.17 mM KCl, 0.33 mM CaCl₂, 0.33 mM MgSO_4_ in distilled water, dH_2_O) and methylene blue (0.5 ppm, M9140; Sigma-Aldrich) as a fungicide, at 28°c. Zebrafish embryos were checked daily, and dead embryos were removed. At 6 dpf, surviving larvae were recorded, and this count was used as the starting population for survival analysis. The 6 dpf larvae were transferred to 3.5 L tanks containing reverse osmosis water (maintained at pH 7.5) at a seeding density of 60 fish per tank. Zebrafish were monitored daily during feeding and counted weekly. As a humane endpoint, moribund zebrafish were euthanised with a tricaine overdose (4 mg/ml) until respiration ceased to confirm death.

### Adipose Staining

Zebrafish larvae were stained in Nile Red (#N1142, Sigma-Aldrich), which labels neutral lipids, as previously described^52,53^. Briefly, zebrafish larvae were immersed in E3 containing 0.5μl/ml Nile red (1.25 mg/mL) for 30 minutes before image acquisition on a Stereomicroscope (NSZ-606 Binocular Zoom) fitted with a Nightsea SFA light base. For confocal microscopy at cellular resolution, larvae were first Nile red stained, euthanised in tricaine and then submerged in a fixative clearing solution (4% Neutral Buffered Formalin, 4% KOH, 1% Triton-X) for 48 hours at 42°C to optimise specimen transparency, as described previously^54^.

### Image Analysis

Images obtained through stereo microscopy were analysed with ImageJ software (version 1.54g, NIH, USA). The image scale was calibrated using a ruler scale bar placed next to the zebrafish samples. The adipose and liver areas were delineated based on the thresholding of pixel intensity. Lateral view images were used for all measurements.

### Chemical Treatment

Zebrafish larvae were raised conventionally until the start of the experiment. For bile salt exposure experiments, larvae were randomly allocated to control or treated groups. The vehicle control group was exposed to 0.1% dimethyl sulfoxide (DMSO, Sigma-Aldrich) in E3 medium. In contrast, treated fish were exposed to 1 mM Taurocholic acid (TCA, #16215, Cayman Chemical) in the presence or absence of 1 µM Odevixibat (Odev, #35344, Cayman Chemical) in E3. The media in each plate was changed daily, and larvae were fed fresh paramecia and brine shrimp. For experiments involving P53KO;lf:YAP zebrafish, larvae were raised until 14 dpf. At this stage, larvae were screened for liver tumour development under a fluorescence microscope. PY-TB fish were randomly assigned to control or treated groups. Vehicle control PY-TB groups received 0.1% DMSO in E3, while treated groups were exposed to 1.5 µM Odevixibat in E3. Media in all plates was changed daily, and larvae were fed fresh paramecia and brine shrimp.

### Histology and Immunofluorescence

Zebrafish samples were fixed in 4% paraformaldehyde (PFA, 441244, Sigma-Aldrich) overnight at 4℃. Samples were arranged in agarose-embedded arrays, then paraffin-embedded and serially sectioned at a thickness of 8 μm. For histological analysis, slides were stained with Haematoxylin and Eosin (H&E). Sections were imaged with an Olympus BX53 microscope. For immunofluorescence analysis, sections were initially unmasked using a citrate buffer antigen protocol, as previously described^55,56^. Slides were subsequently stained with primary antibodies against GFP (chicken, 1:400, #ab13970, Abcam), S6 ribosomal protein (rabbit, 1:300, #2217, Cell Signaling Technology), Phospho-S6 ribosomal protein Ser235/236 (rabbit, 1:300, #4858S, Cell Signaling Technology), and 2F11 (Annexin A4) (mouse, 1:300, #ab71286, Abcam). Primary antibodies were diluted in 5% Bovine Serum Albumin (BSA, A7906, Sigma Aldrich) in PBS, and slides were incubated overnight at 4°c. Slides were washed several times with PBS + 0.1% Tween. Bound antibodies were visualised with secondary antibodies conjugated to Alexa Fluor™ 488 (anti-chicken, 1:250, # A-11039, Thermo Scientific), Highly Cross-Adsorbed Alexa Fluor™ 488 (anti-rabbit, 1:250, # A-11034, Thermo Scientific), Highly Cross-Adsorbed Alexa Fluor™ 647 (anti-mouse, 1:250, # A-31571, Thermo Scientific), Cross-Adsorbed Alexa Fluor™ 647 (anti-rat, 1:250, # A-21247, Thermo Scientific). Sections were imaged with a FLUOVIEW FV3000 confocal scanning microscope (Olympus).

### Bromodeoxyuridine staining and analysis

BrdU staining and imaging were performed as previously described^57^. Briefly, 21 dpf zebrafish were exposed to 5 mg/ml Bromodeoxyuridine (BrdU [B5002, Sigma-Aldrich]) in 0.5% DMSO containing E3 medium for 24 hours. Fish were washed five times for 30 minutes each in E3, before being fixed in 4% paraformaldehyde (PFA) overnight at 4℃. Whole fish were arranged in agarose-embedded arrays, then embedded in paraffin and serially sectioned at 8 μm thickness. Sections were unmasked using a citrate buffer antigen retrieval protocol, as previously described^55^. Sections were equilibrated in a DNAse buffer for 30 minutes at 37℃, then treated with DNAse reaction mix (Direct-zol RNA Miniprep kit, Zymo Research) for 60 minutes at 37℃. Sections were incubated with 5% Bovine serum albumin (BSA) for 2 hours at room temperature (RT). They were then incubated with anti-BrdU (Abcam, #ab6326) at a 1:100 dilution overnight at 4℃. After washing with PBS + 0.1% Tween (PBST), sections were incubated with secondary antibody anti-rat Alexa Fluor 647 (Thermo Scientific) at 1:250, and 4’,6-diamidino-2-phenylindole (DAPI) [D3571, Thermo Scientific] at 1:1000 dilution in BSA for 45 minutes at RT. The sections were washed in PBST, then imaged using a FLUOVIEW FV3000 confocal scanning microscope. Images were processed, analysed, and quantified in ImageJ (version 1.54g, NIH, USA).

### Micro-CT Imaging and Analysis

At 21 dpf, zebrafish were collected and fasted for 6 hours prior to being euthanised with tricaine and fixed overnight in 4% paraformaldehyde at 4°C. The samples were washed in PBS and gradually dehydrated in 35% and 50% ethanol (EtOH), before being stained in 1% phosphotungstic acid (PTA) [P4006, Sigma-Aldrich] in EtOH for 3 days with gentle agitation at room temperature, as previsouly described^58^. After staining, the samples were washed in 70% EtOH, and Micro-CT imaging was performed with a Phoenix Nanotom M (Waygate Technologies) using an 80 kV, 180 μA X-ray beam and a 0.2 mm aluminium filter. PTA-stained samples were mounted in micropipette tips and scanned in 70% EtOH using a diamond-coated tungsten target. Standard scans took 24 minutes, acquiring 1200 projections over 360°, with 0.5 s exposure and 2× averaging. Voxel resolution ranged from 1.5 to 1.7 μm, focusing on a mid-abdominal region of interest. Some whole-body scans used a 2.5 μm resolution at 60 kV, 300 μA. Volume reconstructions were generated with datos|x software, utilising inline median and ROI-CT filters, and exported in 32-bit format for further analysis. Reconstructed data were imported into Avizo3DPro (v2019.3), normalised to 16-bit, and reoriented for sagittal and transverse views. Liver, adipose, and muscle tissues were segmented using a combination of keyframe interpolation, grey-value thresholding, and ambient occlusion to identify low-density porous areas. Segmentation was reviewed independently and refined as necessary. Tissue volumes were quantified using Avizo’s volume fraction tools. Muscle area was assessed at the C8 vertebra cross-section by measuring the proportion of muscle area relative to the entire specimen cross-section.

### RNA-seq and analysis

Whole zebrafish larvae were euthanised in tricaine, pooled into microcentrifuge tubes, snap-frozen in liquid nitrogen, and stored at −80 °C. Total RNA was extracted using the Direct-zol RNA Miniprep kit (R2052, Zymo Research) following homogenisation in TRIzol reagent (15596026, Invitrogen), according to the manufacturer’s protocol. RNA integrity and concentration were assessed with the Agilent TapeStation system. RNA-seq libraries were prepared using the Lexogen QuantSeq 3′ mRNA-Seq protocol and sequenced on an Illumina NextSeq 2000 platform with 100 bp single-end reads. Sequencing depth ranged from 5.9 to 9.9 million reads per sample (mean depth: 8.35 million). Raw sequencing files were converted to FASTQ format using the Galaxy platform^59^. Reads were initially aligned to the danRer10 genome with Seqliner v2.6 and subsequently re-aligned to the GRCz11 zebrafish genome build. Gene-level quantification was performed using the Lawson Lab Zebrafish Transcriptome annotation (version 4.2.1)^60^. Transcriptional signatures were examined using Gene Set Enrichment Analysis (GSEA v4.0.3)^61^.

### Spatial transcriptomics (Stereo-seq) sequencing and analysis

Zebrafish were euthanised using tricaine and embedded in optimal cutting temperature (OCT, [4583, Tissue-Tek]) in tissue moulds, then snap frozen. Frozen tissue blocks were stored at −80 °C and cryosectioned at 10 µm onto STOmics Stereo-seq chips (V1.1; BGI Research). Adjacent sections were stained with H&E. Sections were mounted on the Stereo-seq chips and fixed in methanol at −20℃ for 30min, followed by fluorescent staining using the Qubit ssDNA reagent and imaged at 10X magnification with a Motic Microscope. The sections then underwent enzymatic permeabilization for 20 min to release mRNA for capture by spatially barcoded nanoballs on the chip surface. Reverse transcription was performed for 1h at 42℃ using RT QC Mix, after tissue removal, library construction were performed following the STOmics Stereo-seq protocol (v1.1, BGI Genomics), with spatial barcodes and sequencing adapters incorporated during library preparation, as previously described.^20,21^. Library quality was assessed using a Qubit fluorometer and Agilent Bioanalyzer, and sequencing was conducted on a DNBSEQ-T7 platform (MGI) with paired-end 100 bp reads. Raw data were processed through the Stereo-seq pipeline (SAW v1.0.0) to generate spatially resolved transcript count matrices. Transcriptomic data with square bin (Bin50) were processed in R (v4.4.0) using Seurat (v5.1.0) and ggspavis (v1.10.0). Spots with >30% mitochondrial gene content or fewer than 100 total UMI counts were excluded. Normalisation was performed using SCTransform with a clipping range of −10 to 10. Principal component analysis (PCA) was conducted on the top 30 components and used to generate Uniform Manifold Approximation and Projection (UMAP) plots. Louvain clustering (resolution = 0.5) was applied using Seurat’s FindClusters function. Spatial feature plots and cluster maps were visualised using ggspavis and ggplot2.

### Zebrafish CRISPR/Cas9 gene editing

Alt-R CRISPR-Cas9 crRNAs (Integrated DNA Technologies, IDT) targeting exonic regions of *slc45a2*, *cyp7a1* and *tgr5* were used to perform global editing in zebrafish larvae as previously described^62^. crRNA guides were combined with recombinant Alt-R S.p. Cas9 Nuclease V3 (1081058, IDT) in vitro before microinjection. Cas9-guide complex reactions included 12 μM crRNA site 1, 12 μM crRNA site 2, 24 μM Alt-R tracrRNA (IDT), and 5 μg Cas9 in a total volume of 5 μl. These complexes were injected into one-cell stage zebrafish embryos. Target sites for gene insertion or deletion (indel) with CRISPR/Cas9 in zebrafish are listed below. Gene editing was carried out using 1-2 independent Alt-R CRISPR-Cas9 crRNAs targeting exonic regions of cyp7a1 or tgr5, together with 2 independent crRNAs targeting slc45a2.

#### Target sites

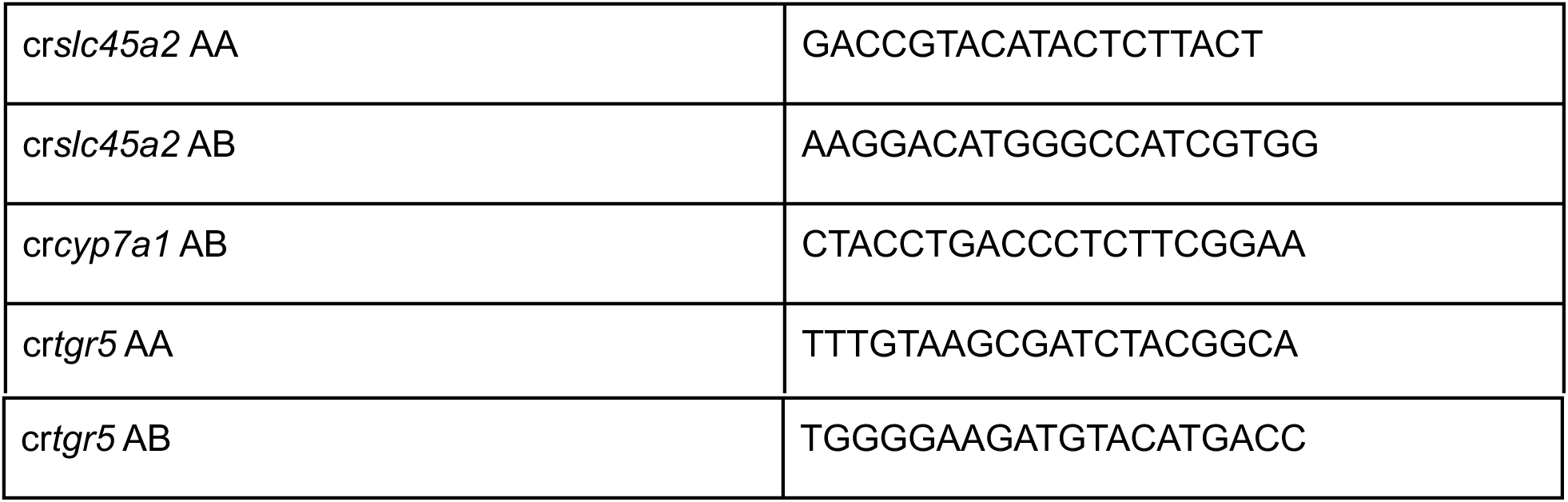

### Gene editing efficiency assessment

Genomic DNA was extracted from injected zebrafish larvae using NB buffer (10 mM Tris-HCl pH 8, 1 mM EDTA, 50 mM KCl, 0.3% TWEEN-20, and 0.3% NP-40) with proteinase K (Thermo Fisher Scientific). PCR was performed using KOD Hot Start Polymerase (Sigma-Aldrich). Sanger sequencing was conducted on gel-purified amplicons by the Australian Genome Research Facility (AGRF). Indels in the sequenced data were analysed using the Inference of CRISPR Edits (ICE) web-based platform (Synthego). Primers used for Sanger sequencing and ICE analysis are shown below.

### Oligonucleotides

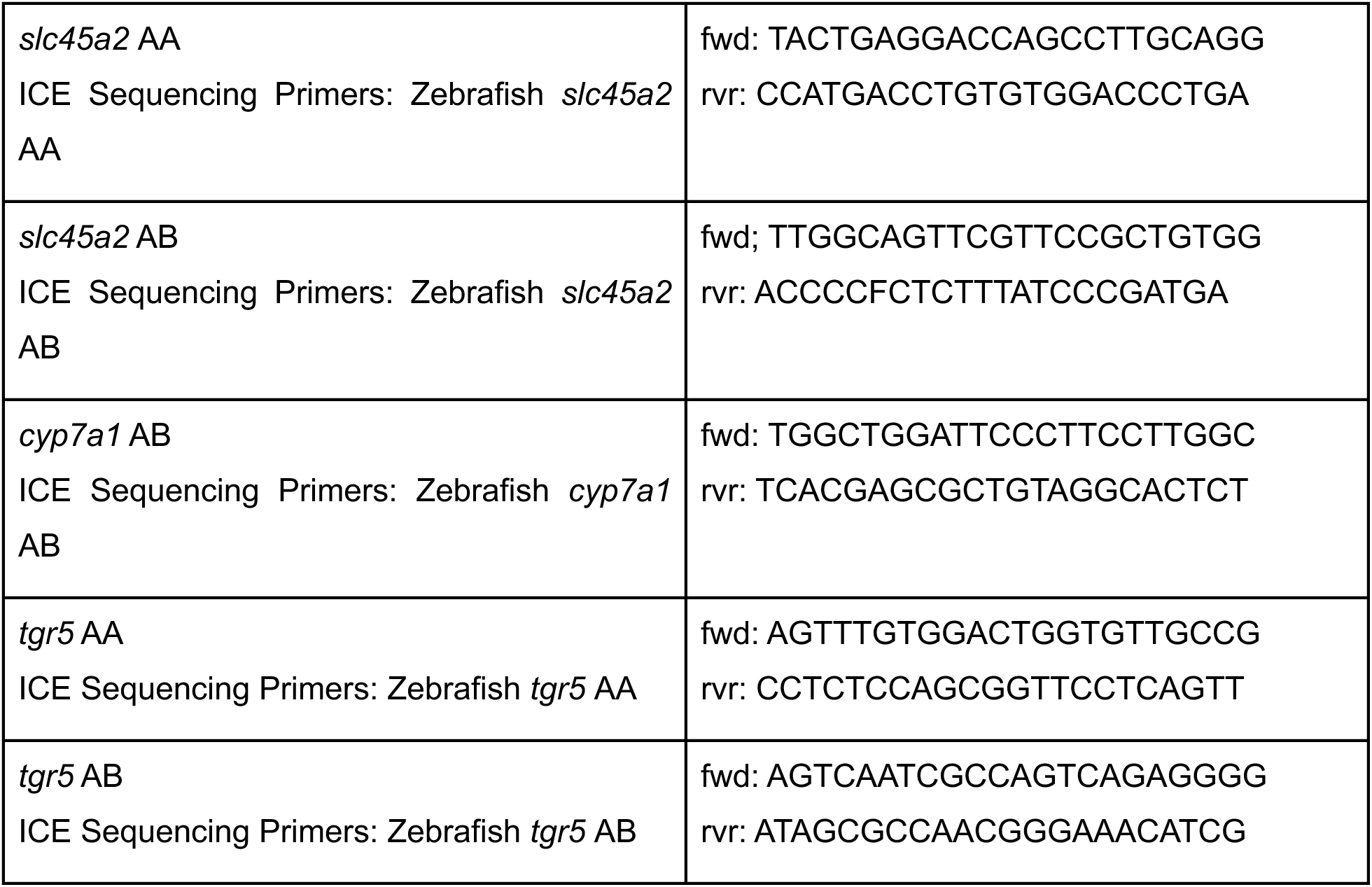

### AAV injection in mice

AAV viruses were diluted in sterile PBS; 1 × 10¹¹ genome copies (GC) of each AAV were administered to 3-week-old mice via retro-orbital injection at a volume of 30 μl. *pAAV-TBG-Cre* (Addgene, #107787) and *pAAV-TBG-Null* (Addgene, #105536) were purchased from Addgene, whereas *pAAV-TBG-YAP^S127A^* and *pAAV-TBG-Bsep* plasmids were generated as previously described^26^. All AAVs were produced in the lab. For YAPS127A overexpression in the TetO-YAP mice, 1 × 10^11 GC of AAV-TBG-Cre were administered to 8-week-old TetO-YAP mice. Two weeks later, YAPS127A expression was induced in the mice by feeding them Dox water (0.2 g/L).

### Quantification and Statistical Analysis

All statistical analyses were performed using Prism v10.0 (GraphPad Software, La Jolla, CA). For comparisons involving more than two groups, datasets were first assessed for normality using the Shapiro–Wilk test. If normality was met, variance homogeneity was tested using Brown–Forsythe’s test. When both assumptions were satisfied, differences were evaluated using one-way ANOVA with Tukey’s multiple-comparisons test. If normality was met but variances were unequal, Welch’s one-way ANOVA with Dunnett’s T3 multiple-comparisons test was used. When normality was not met, Kruskal–Wallis tests with Dunn’s multiple-comparisons correction were applied. For comparisons between two groups, unpaired two-tailed Student’s t-tests were used when assumptions of normality and equal variance were met. When variances were unequal, Welch’s t-test was applied. For non-parametric data, the Mann–Whitney U test was used. Survival analyses were performed using Kaplan–Meier survival curves, and differences between groups were assessed using the log-rank (Mantel–Cox) test. Significance is indicated as *p<0.05, **p<0.01, ***p<0.001 and ****p<0.0001. Unless otherwise stated, data are presented as mean ± s.e.m.

## Acknowledgements

We acknowledge support from the Peter MacCallum Cancer Centre Foundation and the Australian Cancer Research Foundation. We extend our thanks to the Peter MacCallum Cancer Centre Core Facilities and their staff who provided support for this work; namely the Molecular Genomics Core (RRID:SCR_025695), the Centre for Advanced Histology and Microscopy (RRID:SCR_025432), and Research Laboratory Support Services (RRID:SCR_025699). We thank the Melbourne Trace Analysis for Chemical, Earth and Environmental Sciences (TrACEES) Platform for access to the micro-CT scanner, the DrUM Zebrafish Core Facility (University of Melbourne), and BGI Research for assistance with Stereo-seq. We thank the following funders for fellowship, scholarship, and grant support: NHMRC Project Grant 1146558, NHMRC Investigator Grant GNT1176650, NHMRC Ideas Grant 2037181, Peter MacCallum Cancer Foundation, GESA Project Grant, VCA MCRF23013 and a Dame Kate Campbell Fellowship (A.G.C); Peter MacCallum Postgraduate Scholarship (M.W.). The funders had no role in study design, data collection, analysis, or decision to publish or preparation of the manuscript. Lastly, we extend our gratitude to members of the Cox and Brown Laboratories (Peter MacCallum Cancer Centre) for their helpful discussions.

---

**Figure S1.**
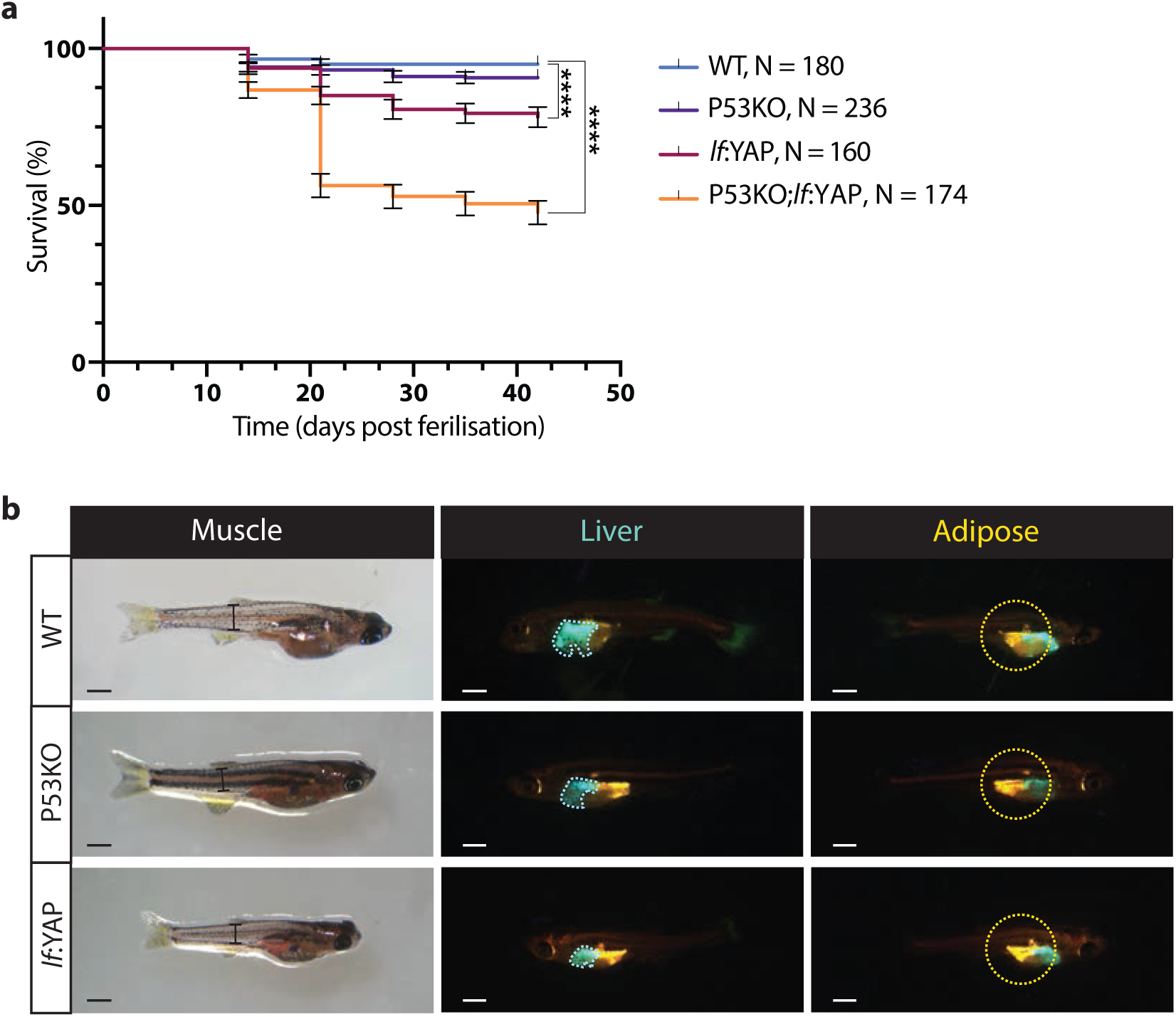
Development of a YAP-driven model of liver cancer that exhibits cachexia. **a**, Kaplan–Meier survival curves of wild-type (WT; n = 180), P53KO (n = 236), *lf*:YAP (n = 160) and P53KO;*lf*:YAP (PY; n = 174) zebrafish. **b**, Representative images of 21 dpf WT, P53KO, lf:YAP and PY zebrafish showing gross morphology (brightfield, left), lf:CFP-NTR liver fluorescence (middle) and Nile Red-stained adipose tissue with liver fluorescence (right). In brightfield images, black lines indicate muscle width measurement. Scale bar, 1 mm. All data are mean ± s.e.m. P values were determined by log-rank (Mantel–Cox) test (a). *P < 0.05, **P < 0.01, ***P < 0.001, ****P < 0.0001.

**Figure S2.**
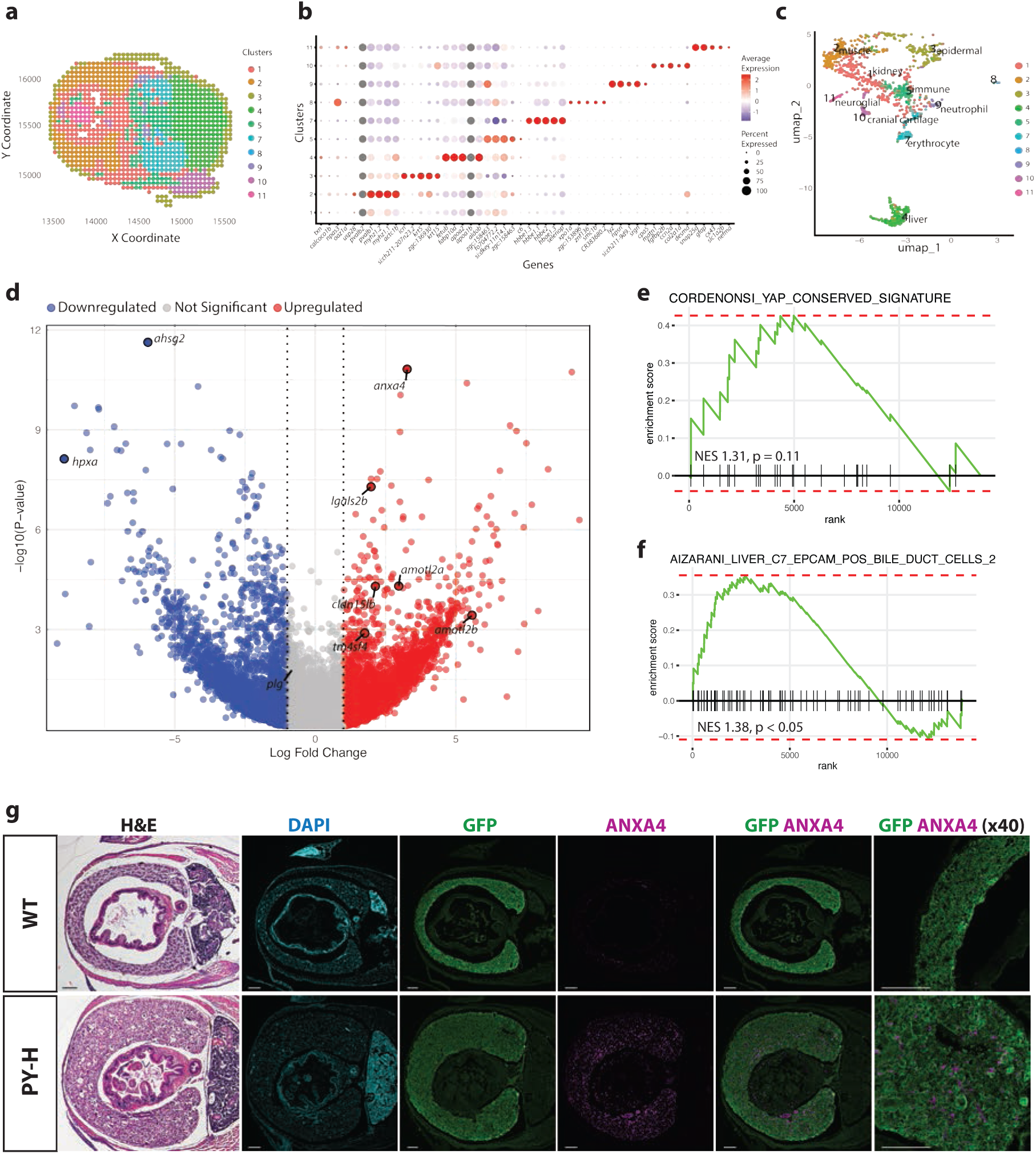
Spatial transcriptomic analysis reveals a signature of liver cancer-associated cachexia. **a**, Spatial feature plot of WT sample showing spot-level clustering derived by principal component analysis and graph-based clustering in Seurat. **b**, Dot plot showing the top three cluster-defining genes per WT cluster, identified by Seurat marker gene analysis. **c**, UMAP of spatial transcriptomic spots from the WT sample coloured by Seurat-defined cluster identity. Cluster annotation was guided by regionally enriched marker genes, with tissue or cell type assignments informed by Zebrahub. **d**, Volcano plot of differentially expressed genes (DEGs) from RNA-seq of dissected livers, comparing P53KO;*lf*:YAP tumour-burdened (PY-TB) fish and WT controls at 21 dpf. Cholangiocyte markers (anxa4, igals2b), hepatocyte markers (ahsg2, hpxa, plg) and YAP target genes (tm4sf4, amotl2b, amotl2a, cldn15lb) are labelled. **e-f**, Gene set enrichment analysis (GSEA) plots of selected pathways enriched in PY-TB versus WT livers, derived from RNA-seq analysis. **g**, H&E and immunofluorescence staining of livers from WT and P53KO;*lf*:YAP hyperplastic (PY-H) fish at 21 dpf. Nuclei are marked with DAPI (cyan), hepatocytes with GFP (green), and cholangiocytes with ANXA4 (magenta). Scale bar, 100 μm.

**Figure S3.**
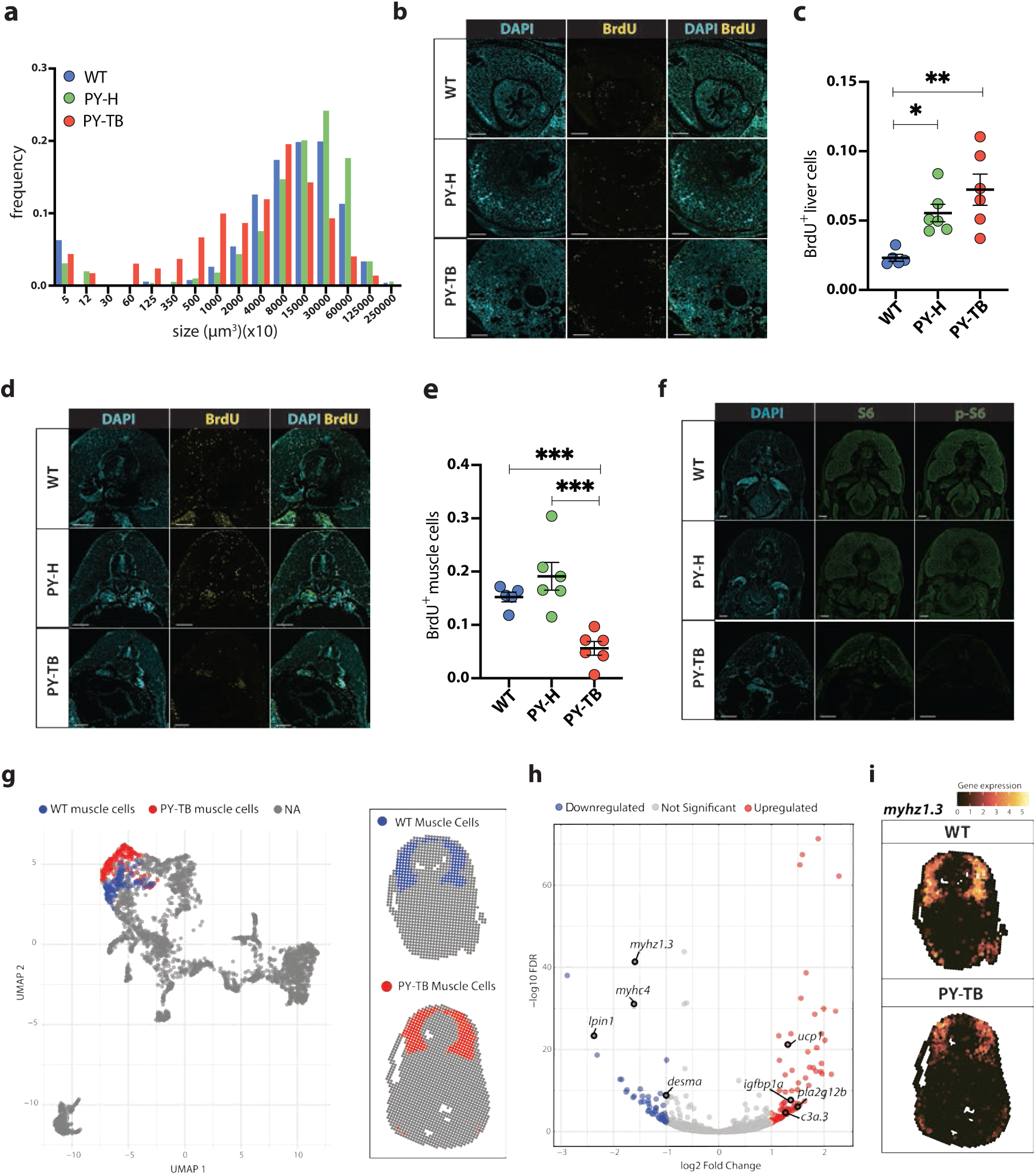
YAP disrupts bile acid homeostasis and drives muscle and adipose wasting. **a**, Histogram of relative adipocyte volume distributions in WT, PY-H and PY-TB zebrafish at 21 dpf. Nile Red–stained adipocytes were imaged by FV300 confocal microscopy and segmented using Imaris. Distributions were normalized within each fish group to allow comparison of adipocyte size independent of adipocyte number. **b**, Representative confocal images of BrdU incorporation (yellow) in liver tissue from WT, PY-H, and PY-TB zebrafish at 21 dpf. **c,** Quantification of BrdU incorporation in liver, expressed as the ratio of BrdU⁺ cells relative to total DAPI⁺ nuclei. **d,** Representative confocal images of BrdU incorporation (yellow) in muscle tissues from WT, PY-H and PY-TB zebrafish at 21 dpf. **e,** Quantification of BrdU incorporation in muscle expressed as the ratio of BrdU⁺ cells relative to total DAPI⁺ nuclei. **f,** Representative confocal images of phospho-S6 and total S6 antibody staining in skeletal muscle of WT, PY-H, and PY-TB zebrafish at 21 dpf. **g,** UMAP and spatial feature plots of muscle-enriched cells identified from spatially resolved transcriptomics (SRT) using a Seurat-based clustering pipeline. Cells were selected from a muscle-associated cluster. WT muscle cells, blue; PY-TB muscle cells, red. **h,** Volcano plot of differentially expressed genes (DEGs) between PY-TB and WT muscle cells from SRT data. **i**, Spatial projection of myogenic marker gene, *myhz1.3* expression over the tissue section from SRT. All scale bars, 100 μm. All data are mean ± s.e.m. P values were determined by one-way ANOVA with Tukey’s multiple-comparisons test (**g,h**). *P < 0.05, **P < 0.01, ***P < 0.001, ****P < 0.0001.

**Figure S4.**
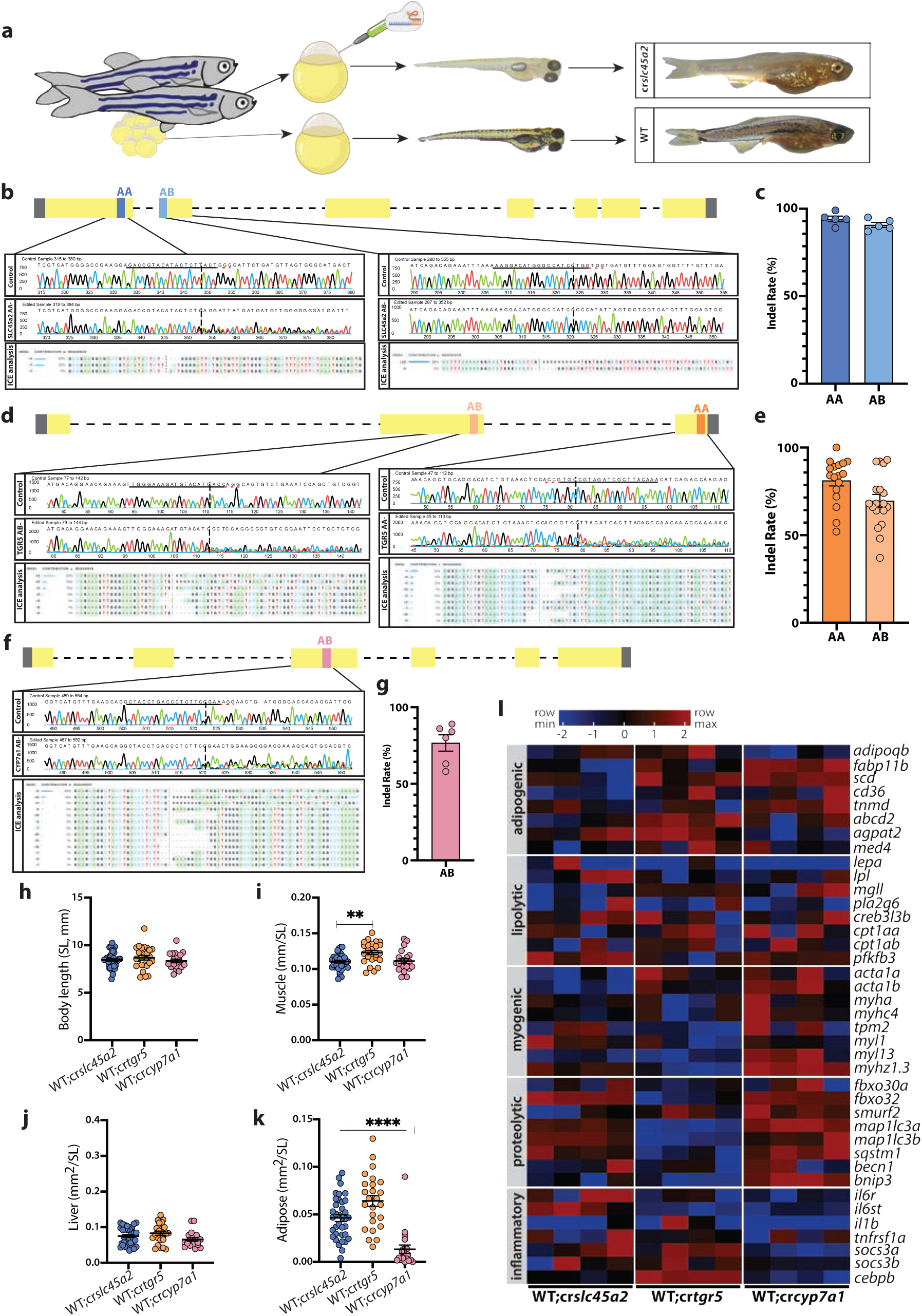
YAP-induced cachexia requires CYP7A1 and TGR5. **a**, Schematic of the CRISPR–Cas9 knockout strategy targeting *slc45a2*, designed to induce a visible pigmentation-loss phenotype for screening. Representative brightfield images of 6 and 21 dpf WT and cr*slc45a2* zebrafish show loss of melanin pigmentation in mutants. **b**, Schematic of CRISPR guide RNA target sites (aa and ab) on the *slc45a2* gene with representative Sanger sequencing traces and ICE analysis of indels from 7 dpf larvae injected at the one-cell stage with both guides. **c**, Indel efficiency rates for each *slc45a2* CRISPR target site. **d**, Schematic of CRISPR guide RNA target sites (aa and ab) on the *tgr5* gene with representative Sanger sequencing traces and ICE analysis from 7 dpf larvae co-injected at the one-cell stage with *tgr5* and *slc45a2* guides (latter not shown). **e**, Indel efficiency rates for each *tgr5* CRISPR target site. **f**, Schematic of the CRISPR target site (ab) on *cyp7a1* with corresponding Sanger sequencing traces and ICE analysis from 7 dpf larvae co-injected at the one-cell stage with *cyp7a1* and *slc45a2* guides (latter not shown). **g**, Indel efficiency rates for the *cyp7a1* CRISPR target site. **h-k**, Biometrics of body length (**h**, standard length, SL), muscle width (**i**), liver size (**j**) and adipose tissue size (**k**) in WT cr*slc45a2*, WT cr*tgr5* and WT cr*cyp7a1* zebrafish at 21 dpf. **l**, Heat map of selected adipogenic, lipolytic, myogenic, proteolytic and inflammatory gene expression profiles in pooled WT, *crslc45a2*, *crtgr5* and *crcyp7a1* zebrafish at 21 dpf, determined by bulk RNA-seq (n = 4 samples of 5 fish per condition). All data are mean ± s.e.m. P values were determined by one-way ANOVA with Tukey’s multiple-comparisons test (**h,i**) and Kruskal–Wallis test with Dunn’s multiple-comparisons post hoc adjustment (**j,k**). *P < 0.05, **P < 0.01, ***P < 0.001, ****P < 0.0001.

**Figure S5.**
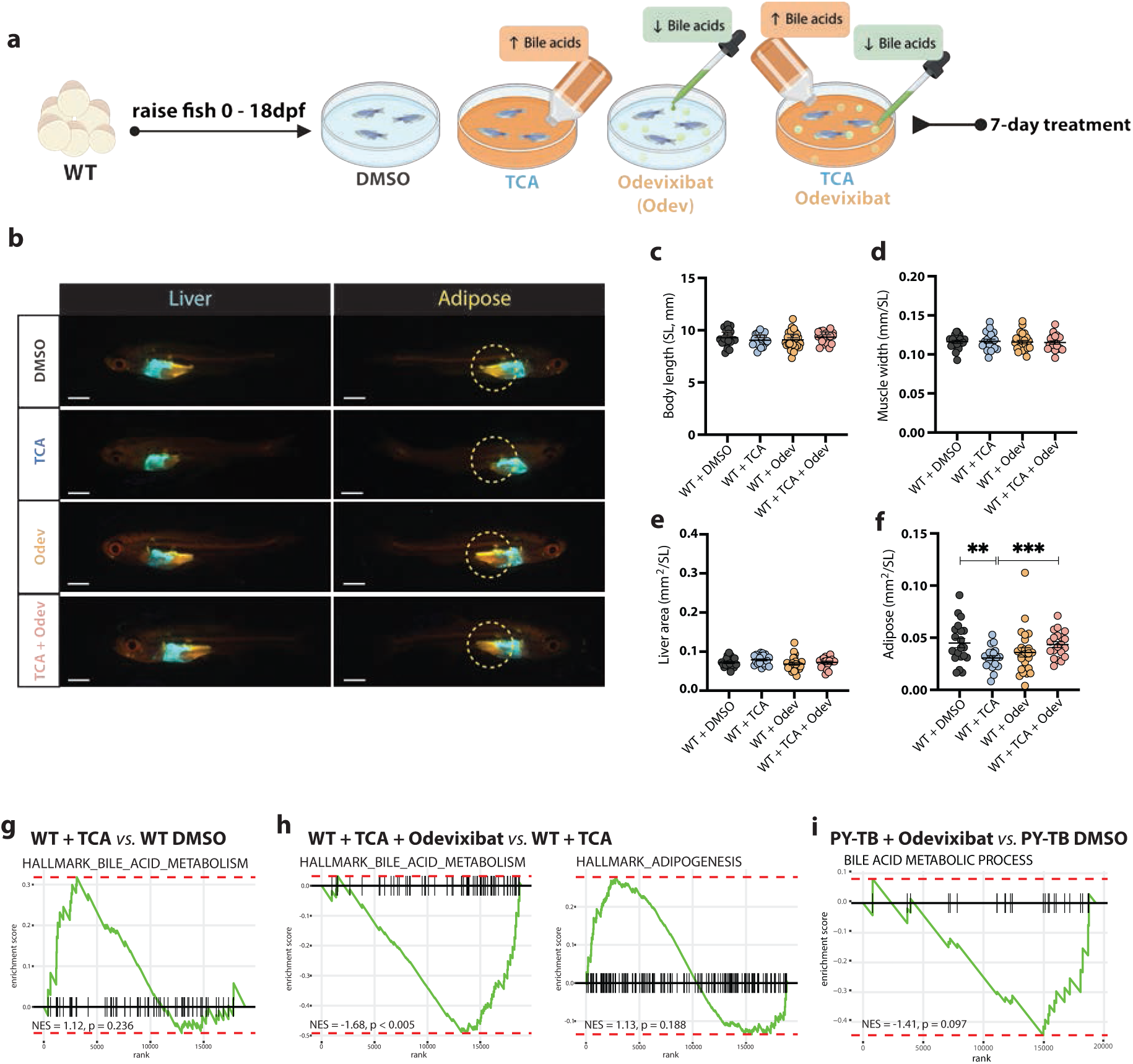
Odevixibat restores bile acid homeostasis and prevents cancer-associated cachexia. **a**, Schematic of the bile acid (taurocholic acid, TCA) and Odevixibat (Odev) treatment intervention protocol. **b**, Representative images of 25 dpf WT zebrafish following DMSO (control), TCA, Odev, or combinatorial TCA + Odev tretament. Liver tissue, *lf*:CFP-NTR fluorescence (cyan); adipose tissue, Nile Red staining (yellow). Scale bar, 1 mm. **c-f**, Biometrics of body length (**c**, standard length, SL), muscle width (**d**), liver size (**e**) and adipose tissue size (**f**) in WT zebrafish across the four treatment groups at 25 dpf. **g**, GSEA of bulk RNA-seq comparing TCA-treated WT zebrafish versus DMSO-treated WT zebrafish at 25 dpf. Bulk RNA-seq was performed on 4 pooled samples of 5 fish per condition. **h**, GSEA of bulk RNA-seq comparing combined TCA and Odev treatment versus TCA treatment in WT zebrafish at 25 dpf. Bulk RNA-seq was performed on 4 pooled samples of 5 fish per condition. **i**, GSEA of bulk RNA-seq comparing PY-TB Odev-treated zebrafish versus PY-TB DMSO-treated control counterparts at 21 dpf following treatment. Bulk RNA-seq was performed on 4 pooled samples of 5 fish per condition. All data are mean ± s.e.m. P values were determined by one-way ANOVA with Tukey’s multiple-comparisons test (**c,d**) and Kruskal–Wallis test with Dunn’s multiple-comparisons post hoc adjustment (**e,f**). *P < 0.05, **P < 0.01, ***P < 0.001, ****P < 0.0001.

## References

1. Ferrer, M. et al. Cachexia: A systemic consequence of progressive, unresolved disease. Cell 186, 1824–1845 (2023).

2. Swanton, C. et al. Embracing cancer complexity: Hallmarks of systemic disease. Cell 187, 1589–1616 (2024).

3. Babic, A. et al. Adipose tissue and skeletal muscle wasting precede clinical diagnosis of pancreatic cancer. Nat. Commun. 14, 4317 (2023).

4. Al-Sawaf, O. et al. Body composition and lung cancer-associated cachexia in TRACERx. Nat. Med. 29, 846–858 (2023).

5. Groarke, J. D. et al. Ponsegromab for the treatment of cancer cachexia. N. Engl. J. Med. 391, 2291–2303 (2024).

6. Harvey, K. F., Zhang, X. & Thomas, D. M. The Hippo pathway and human cancer. Nat. Rev. Cancer 13, 246–257 (2013).

7. Ma, S., Meng, Z., Chen, R. & Guan, K.-L. The Hippo Pathway: Biology and Pathophysiology. Annu. Rev. Biochem. 88, 577–604 (2019).

8. Yimlamai, D. et al. Hippo pathway activity influences liver cell fate. Cell 157, 1324–1338 (2014).

9. Sugihara, T., Isomoto, H., Gores, G. & Smoot, R. YAP and the Hippo pathway in cholangiocarcinoma. J. Gastroenterol. 54, 485–491 (2019).

10. Banales, J. M. et al. Cholangiocarcinoma 2020: the next horizon in mechanisms and management. Nat. Rev. Gastroenterol. Hepatol. 17, 557–588 (2020).

11. Cox, A. G. et al. Yap reprograms glutamine metabolism to increase nucleotide biosynthesis and enable liver growth. Nat. Cell Biol. 18, 886–896 (2016).

12. Cox, A. G. et al. Yap regulates glucose utilization and sustains nucleotide synthesis to enable organ growth. EMBO J. 37, (2018).

13. Vaidyanathan, S. et al. YAP regulates an SGK1/mTORC1/SREBP-dependent lipogenic program to support proliferation and tissue growth. Dev. Cell 57, 719–731.e8 (2022).

14. Ibar, C. & Irvine, K. D. Integration of Hippo-YAP Signaling with Metabolism. Dev. Cell 54, 256–267 (2020).

15. Kobayashi, S., Cox, A. G., Harvey, K. F. & Hogan, B. M. Vasculature is getting Hip(po): Hippo signaling in vascular development and disease. Dev. Cell 58, 2627–2640 (2023).

16. Ali, E. S. & Ben-Sahra, I. Regulation of nucleotide metabolism in cancers and immune disorders. Trends Cell Biol. 33, 950–966 (2023).

17. Moya, I. M. et al. Peritumoral activation of the Hippo pathway egectors YAP and TAZ suppresses liver cancer in mice. Science 366, 1029–1034 (2019).

18. Algueró-Nadal, A. et al. Steatotic liver disease induces YAP/TAZ-driven cell competition that can suppress tumor initiation. J. Hepatol. (2025) doi:10.1016/j.jhep.2025.06.002.

19. Berghmans, S. et al. tp53 mutant zebrafish develop malignant peripheral nerve sheath tumors. Proc. Natl. Acad. Sci. U. S. A. 102, 407–412 (2005).

20. Liu, C. et al. Spatiotemporal mapping of gene expression landscapes and developmental trajectories during zebrafish embryogenesis. Dev. Cell 57, 1284–1298.e5 (2022).

21. Chen, A. et al. Spatiotemporal transcriptomic atlas of mouse organogenesis using DNA nanoball-patterned arrays. Cell 185, 1777–1792.e21 (2022).

22. Sur, A. et al. Single-cell analysis of shared signatures and transcriptional diversity during zebrafish development. Dev. Cell 58, 3028–3047.e12 (2023).

23. Wen, J. et al. Fxr signaling and microbial metabolism of bile salts in the zebrafish intestine. Sci. Adv. 7, eabg1371 (2021).

24. Thompson, R. J. et al. Odevixibat treatment in progressive familial intrahepatic cholestasis: a randomised, placebo-controlled, phase 3 trial. Lancet Gastroenterol Hepatol 7, 830–842 (2022).

25. Ray, K. Positive phase III results for odevixibat for progressive familial intrahepatic cholestasis. Nat. Rev. Gastroenterol. Hepatol. 19, 556 (2022).

26. Liu, Y. et al. Disrupting bile acid metabolism by suppressing Fxr causes hepatocellular carcinoma induced by YAP activation. Nat. Commun. 16, 3583 (2025).

27. Kwon, Y. et al. Systemic organ wasting induced by localized expression of the secreted insulin/IGF antagonist ImpL2. Dev. Cell 33, 36–46 (2015).

28. Figueroa-Clarevega, A. & Bilder, D. Malignant Drosophila tumors interrupt insulin signaling to induce cachexia-like wasting. Dev. Cell 33, 47–55 (2015).

29. Stacchiotti, S. et al. GDF-15 predicts epithelioid hemangioendothelioma aggressiveness and is downregulated by sirolimus through ATF4/ATF5 suppression. Clin. Cancer Res. 30, 5122–5137 (2024).

30. Marti, P. et al. YAP promotes proliferation, chemoresistance, and angiogenesis in human cholangiocarcinoma through TEAD transcription factors. Hepatology 62, 1497–1510 (2015).

31. Chang, L. et al. The SWI/SNF complex is a mechanoregulated inhibitor of YAP and TAZ. Nature 563, 265–269 (2018).

32. Liu, Y. et al. Yap-Sox9 signaling determines hepatocyte plasticity and lineage-specific hepatocarcinogenesis. J. Hepatol. 76, 652–664 (2022).

33. Zhou, Z.-J. et al. Whole-exome sequencing reveals genomic landscape of intrahepatic cholangiocarcinoma and identifies SAV1 as a potential driver. Nat. Commun. 15, 9960 (2024).

34. Ilyas, S. I. et al. Cholangiocarcinoma - novel biological insights and therapeutic strategies. Nat. Rev. Clin. Oncol. 20, 470–486 (2023).

35. Jia, X. et al. Characterization of Gut Microbiota, Bile Acid Metabolism, and Cytokines in Intrahepatic Cholangiocarcinoma. Hepatology 71, 893–906 (2020).

36. Vijay, V. et al. Generation of a biliary tract cancer cell line atlas identifies molecular subtypes and therapeutic targets. Cancer Discov. (2025) doi:10.1158/2159-8290.CD-24-1383.

37. Watanabe, M. et al. Bile acids induce energy expenditure by promoting intracellular thyroid hormone activation. Nature 439, 484–489 (2006).

38. Lee, J. M. et al. Nutrient-sensing nuclear receptors coordinate autophagy. Nature 516, 112–115 (2014).

39. Seok, S. et al. Transcriptional regulation of autophagy by an FXR-CREB axis. Nature 516, 108–111 (2014).

40. Jia, W., Xie, G. & Jia, W. Bile acid-microbiota crosstalk in gastrointestinal inflammation and carcinogenesis. Nat. Rev. Gastroenterol. Hepatol. 15, 111–128 (2018).

41. Velazquez-Villegas, L. A. et al. TGR5 signalling promotes mitochondrial fission and beige remodelling of white adipose tissue. Nat. Commun. 9, 245 (2018).

42. Anakk, S. et al. Bile acids activate YAP to promote liver carcinogenesis. Cell Rep. 5, 1060–1069 (2013).

43. Pepe-Mooney, B. J. et al. Single-cell analysis of the liver epithelium reveals dynamic heterogeneity and an essential role for YAP in homeostasis and regeneration. Cell Stem Cell 25, 23–38.e8 (2019).

44. Ji, S. et al. FGF15 activates Hippo signaling to suppress bile acid metabolism and liver tumorigenesis. Dev. Cell 48, 460–474.e9 (2019).

45. Michalopoulos, G. K. Liver regeneration. J. Cell. Physiol. 213, 286–300 (2007).

46. Merrell, A. J. & Stanger, B. Z. A feedback loop controlling organ size. Developmental cell vol. 48 425–426 (2019).

47. Hanahan, D. & Weinberg, R. A. Hallmarks of cancer: the next generation. Cell 144, 646–674 (2011).

48. Thomas, C., Pellicciari, R., Pruzanski, M., Auwerx, J. & Schoonjans, K. Targeting bile-acid signalling for metabolic diseases. Nat. Rev. Drug Discov. 7, 678–693 (2008).

49. Hagenbeek, T. J. et al. An allosteric pan-TEAD inhibitor blocks oncogenic YAP/TAZ signaling and overcomes KRAS G12C inhibitor resistance. Nat Cancer 4, 812–828 (2023).

50. Chapeau, E. A. et al. Direct and selective pharmacological disruption of the YAP-TEAD interface by IAG933 inhibits Hippo-dependent and RAS-MAPK-altered cancers. Nat. Cancer 5, 1102–1120 (2024).

51. Choi, T.-Y., Ninov, N., Stainier, D. Y. R. & Shin, D. Extensive conversion of hepatic biliary epithelial cells to hepatocytes after near total loss of hepatocytes in zebrafish. Gastroenterology 146, 776–788 (2014).

52. Minchin, J. E. N. & Rawl s, J. F. In vivo analysis of white adipose tissue in zebrafish. Methods Cell Biol. 105, 63–86 (2011).

53. Khan, N. et al. Optimised methods to image hepatic lipid droplets in zebrafish larvae. Dis. Model. Mech. dmm. 050786 (2024).

54. Sakata-Haga, H. et al. A rapid and nondestructive protocol for whole-mount bone staining of small fish and Xenopus. Sci. Rep. 8, 7453 (2018).

55. Brown, R. W. & Chirala, R. Utility of microwave-citrate antigen retrieval in diagnostic immunohistochemistry. Mod. Pathol. 8, 515–520 (1995).

56. Sande-Melon, M., Bergemann, D., Fernández-Lajarín, M., González-Rosa, J. M. & Cox, A. G. Development of a hepatic cryoinjury model to study liver regeneration. Development (2024) doi:10.1242/dev.203124.

57. Tan, V. W. T. et al. SLAM-ITseq identifies that Nrf2 induces liver regeneration through the pentose phosphate pathway. Dev. Cell (2024) doi:10.1016/j.devcel.2024.01.024.

58. Ding, Y. et al. Computational 3D histological phenotyping of whole zebrafish by X-ray histotomography. Elife 8, (2019).

59. Afgan, E. et al. The Galaxy platform for accessible, reproducible and collaborative biomedical analyses: 2016 update. Nucleic Acids Res. 44, W3–W10 (2016).

60. Lawson, N. D. et al. An improved zebrafish transcriptome annotation for sensitive and comprehensive detection of cell type-specific genes. Elife 9, (2020).

61. Subramanian, A. et al. Gene set enrichment analysis: a knowledge-based approach for interpreting genome-wide expression profiles. Proc. Natl. Acad. Sci. U. S. A. 102, 15545–15550 (2005).

62. Ong, A. J. S. et al. The KEAP1-NRF2 pathway regulates TFEB/TFE3-dependent lysosomal biogenesis. Proc. Natl. Acad. Sci. U. S. A. 120, e2217425120 (2023).

